# The bounded rationality of probability distortion

**DOI:** 10.1101/662429

**Authors:** Hang Zhang, Xiangjuan Ren, Laurence T. Maloney

## Abstract

In decision-making under risk (DMR) participants’ choices are based on probability values systematically different from those that are objectively correct. Similar systematic distortions are found in tasks involving relative frequency judgments (JRF). These distortions limit performance in a wide variety of tasks and an evident question is, why do we systematically fail in our use of probability and relative frequency information?

We propose a Bounded Log-Odds Model (BLO) of probability and relative frequency distortion based on three assumptions: (1) *log-odds*: probability and relative frequency are mapped to an internal log-odds scale, (2) *boundedness*: the range of representations of probability and relative frequency are bounded and the bounds change dynamically with task, and (3) *variance compensation*: the mapping compensates in part for uncertainty in probability and relative frequency values.

We compared human performance in both DMR and JRF tasks to the predictions of the BLO model as well as eleven alternative models each missing one or more of the underlying BLO assumptions (factorial model comparison). The BLO model and its assumptions proved to be superior to any of the alternatives. In a separate analysis, we found that BLO accounts for individual participants’ data better than any previous model in the DMR literature.

We also found that, subject to the boundedness limitation, participants’ choice of distortion approximately maximized the mutual information between objective task-relevant values and internal values, a form of bounded rationality.

**Significance Statement:** People distort probability in decision under risk and many other tasks. These distortions can be large, leading us to make markedly suboptimal decisions. There is no agreement on why we distort probability. Distortion changes systematically with task, hinting that distortions are dynamic compensations for some intrinsic “bound” on working memory. We first develop a model of the bound and the compensation process and then report an experiment showing that the model accounts for individual human performance in decision under risk and relative frequency judgments. Last, we show that the particular compensation in each experimental condition serve to maximize the mutual information between objective decision variables and their internal representations. We distort probability to compensate for our own working memory limitations.

---

In making decisions, we choose among actions whose outcomes are typically uncertain; we can model such choices as choices among lotteries. To specify a lottery *L* we list all of its possible outcomes *O*_1_,…,*O*_*n*_ and the corresponding probabilities of occurrence *p*_1_,…,*p*_*n*_ that a specific lottery assigns to each outcome. If we knew all the relevant probabilities, we would be engaged in decision under risk (1). If we can also assign a numerical measure of utility *U* (*O*_*i*_) to each outcome *O*_*i*_, we could assign an *expected utility* to each lottery,

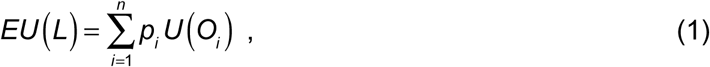

and a decision maker maximizing expected utility (2, 3) would select the lottery with the highest expected utility among those offered. The probabilities serve to weight the contribution of the utility of each outcome. The Expected Utility Theory (EUT) model is simple, but has a wide range of applications, not just in economic decisions but also in perception (4, 5) and planning of movement (6-10).

For more than two centuries EUT was treated as an adequate description of human choice behavior in decision under risk until challenged by Allais (11). In an elegant series of experiments, he showed that human decision makers did not weight utilities by the corresponding probabilities of occurrence in choosing among lotteries. In Prospect Theory, Kahneman and Tversky (12) resolved the Allais paradoxes and other shortcomings of EUT by assuming that decision makers use a transformation of probability *π* (*p*)—a *probability weight* or *decision weight—*in place of probability *p* in the computation of expected utility. The distortion function in decision under risk *π* (*p*) was originally inferred from human choices in experiments and it is often—but not always—an inverted-*S*-shaped function of *p* (13-15).

Wu, Delgado, and Maloney (16) compared performance in a “classical” decision under risk task with performance in a mathematically equivalent motor decision task. Each participant completed both tasks and while the fitted probability distortion functions for the classical task were—as expected—inverted-*S*-shaped, those based on the motor task tended to be better fit by *S*-shaped functions. The same participant could have both the inverted-*S*-shaped and *S*-shaped forms of the distortion function *π* (*p*) in different decision tasks.

Ungemach, Stewart and Chater (17) found a similar tendency to underweight small probabilities in decisions and overweight large (see also 18, 19, 20). Probability distortion in the form of inverted-*S*-shaped and *S*-shaped weighting functions is also found in monkeys’ choice behavior (21) and is supported by human neuroimaging evidence (22, 23).

Zhang and Maloney (24) reported that both the inverted-*S*-shaped or *S*-shaped distortion functions are found in relative frequency and confidence tasks other than decision-making under risk. For convenience, we will use the term “probability” to include relative frequency and confidence. The same participants had different inverted-*S*-shaped or *S*-shaped probability distortion functions in different experimental conditions even though the trials for the different conditions were randomly interleaved. They concluded that the probability distortion function is not fixed for a participant but *dynamic*, changing systematically with task. There is increasing evidence that dynamic remapping of representational range occurs along more abstract dimensions, such as value (25-29), numerosity (30, 31), relative frequency (32), and variance (33).

Zhang and Maloney (24) found that probability distortions could be well fit by the linear transformation:

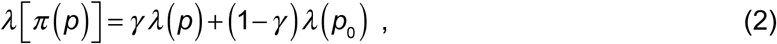

where 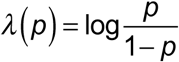 is the log-odds (34) or logit function (35) and *γ* > 0 and 0 < *p*_0_ < 1 are free parameters. See Figure 1a for examples and Zhang and Maloney (24) for further examples, which include 20 datasets taken from 12 studies involving probability, relative frequency and confidence, all the studies for which we could recover and analyze data. We caution that these Linear in Log-Odds (LLO) fits to data represent empirical regularities unmotivated by any theory.

**Figure 1.**
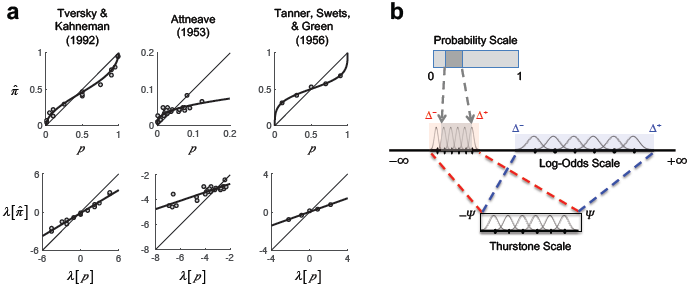
Motivations and intuitions for the bounded log-odds model (BLO). **a**. Observed probability distortions (top row) can be well captured by a linear fit on the log-odds scale (bottom row). The *λ* [*p*] and 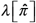 respectively denote the log-odds of the objective and subjective probabilities, *p* and 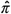. Circles denote data. Thick curves or lines denote the LLO fits. Tversky and Kahneman (1992): Subjective probability (decision weight) versus objective probability in decision under risk. Attneave (1953): Estimated relative frequency of letters in written English versus actual relative frequency. Tanner, Swets, & Green (1956), c.f. Green and Swets (1966/1974): Estimated probability of signal present versus objective probability in a signal detection task. Adapted from (24). **b**. Encoding on the Thurstone scale. A selected range of [Δ^−^, Δ^+^] is encoded on the Thurstone scale [−Ψ,Ψ] with limited resolution. The smaller the encoded range, the smaller the encoding variance.

Over the course of this article we will replace Eq. 2 by a new model, Bounded Log-Odds (BLO) based on theoretical considerations. We propose that probability distortion in both decision under risk and in judgment of relative frequency is fundamentally a consequence of a specific limitation on the dynamic range of the neural representation of probability which we identify. As a consequence of this limitation, human performance in a wide variety of tasks (e.g. the Allais Paradoxes (11)) is necessarily sub-optimal by whatever measure is appropriate to each task.

BLO is based on three assumptions:

1. **log-odds representation**
2. **dynamic encoding on a bounded Thurstone scale**
3. **variance compensation**

We describe these assumptions and possible alternatives in detail below.

We will use factorial model comparison (36) to separately test each of the three assumptions against plausible alternatives. In addition to BLO, we consider eleven variant models each with one or more of the assumptions altered. Half the variant models will have bounded Thurstonian scales, half will not; half will have variance compensation, half will not. We consider two alternatives to the assumption of log-odds representation, giving a total of 2 x 2 x 3 = 12 models, one of which is BLO and one of which is LLO (Eq. 2). We compare human performance to the predictions of each variant model in both a decision-making under risk (DMR) task and also a judgment of relative frequency (JRF) task. Each subject completed both tasks allowing us to compare performance within task.

We will separately compare the performance of BLO to all previous models of decision under risk currently in the literature. The data used in all model comparisons are taken from the new DMR and JRF experiment with 75 participants that we report here and data from a previous article by Gonzalez and Wu (14). We will identify the cognitive constraints in individuals’ representation of probability as well as the optimality under these constraints.

## Maximizing mutual information

The results of our experiments and analyses will indicate that BLO is an accurate *descriptive* model of what participants do in two very different kinds of experiments, DMR and JRF. However, nothing in these analyses serves to demonstrate that BLO is in any sense a normative model or that human performance is normative. In the second part of the article we consider the possibility that the BLO mapping and human performance serve to maximize the mutual information between external decision variables and their internal representation, a form of *bounded rationality* in Herbert Simon’s sense (37). In the last part of the article we show that BLO accounts for a variety of phenomena in DMR.

## Results

### Assumptions of BLO

#### Assumption 1: log-odds representation

In the BLO model probability, *p*, is internally represented as a linear transformation of log-odds,

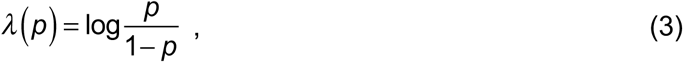

a one-to-one, increasing transformation of probability. A similar log-odds scale has been introduced by Erev and colleagues (38, 39) to explain probability distortion in confidence ratings.

#### Assumption 2: dynamic encoding on a bounded Thurstone scale

Thurstone (40) proposed several alternative models for representing subjective scales and methods for fitting a wide variety of data to such models. We are not concerned with methods for fitting data to Thurstone scales or their use in constructing attitude scales; we are only interested in Thurstone scales as convenient mathematical structures. We can think of the bounded Thurstone scale (40) as an imperfect neural device capable of storing magnitudes within a fixed range. We can encode a magnitude signal *s* anywhere in this range and later retrieve it. The retrieved value *s*, however, is perturbed by Gaussian noise with mean 0 and variance *σ* ^2^ : we might store 0.5 and retrieve 0.63 or 0.48. The schematic Gaussian distributions in Figure 1b capture this representational uncertainty. For simplicity we assume that Gaussian error is independent, identically-distributed across the scale (Thurstone’s Case V).

We could use the entire Thurstonian scale range to represent probabilities from 0 to 1 but—at least in some tasks—only a limited range is needed. For example, in the letter-frequency task of Attneave (41) the probabilities range from about 0.13 (e) to 0.0013 (z) and only a fraction of the full probability scale is needed to carry out the task.

We can pick any interval on the log-odds scale and map it linearly to the Thurstone device. In Figure 1b we illustrate two choices. One represents a small range of the log-odds scale using the full range of the Thurstone device, the other represents a larger range also mapped to the full range of the Thurstone device. The row of Gaussians on the two intervals of the log-odds scale symbolize the encoding uncertainty induced by the Thurstone scale.

The greater the log-odds range that needs to be encoded, the greater the density of the magnitudes along the Thurstone scale, and the greater the chances of confusion of nearby codes and vice versa. The challenge is to choose a transformation that maximizes the information encoded by the scale, which is a problem of ***efficient encoding***. There is experimental evidence for efficient coding in perception (42-45) and recently in perceiving value and probability (27, 29, 46). See especially the review by Simoncelli and Olshausen (44).

Our concern is with the representation of probability, specifically in the form of log-odds. In mathematical notation we select an interval [Δ^−^, Δ^+^] on the log-odds scale to be mapped to the full range of the Thurstone scale [−Ψ,Ψ] and in effect we confine the representation of log-odds *λ* to this interval:

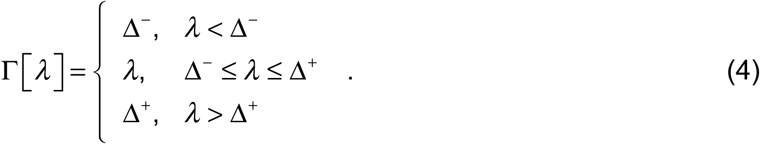

Following the linear mapping from [Δ^−^, Δ^+^] to [−Ψ,Ψ], we have Γ(*λ* [*p*]) mapped to

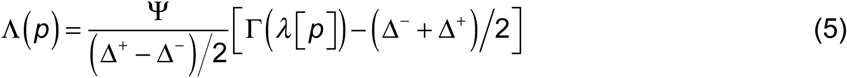

on the Thurstone scale. The neural encoding of *p* can thus be modeled as a Gaussian random variable centered at Λ(*p*), denoted 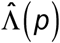. We refer to 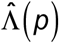 as “truncated log-odds”.

#### Assumption 3: variance compensation

The subjective estimate of probability needed by explicit report or internal use will be decoded from the truncated log-odds encoded on the Thurstone scale. We introduce the transformation

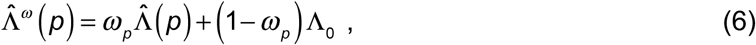

to compensate for encoding uncertainty (see Supplements S1 & S2), where 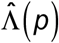 is, as before, the truncated log-odds, Λ_0_ is an anchor point and 0 < *ω* _*p*_ ≤ 1 is a reliability measure of encoding (i.e. inversely related to the variance of encoding) that may vary with *p* (see Methods). The final estimate of probability is 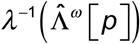, where *λ* ^−1^ (·) denotes the inverse of the logit function.

Similar variance compensation has been widely used to model systematic biases in perception and memory (45, 47). Even probability distortion in the form of LLO is considered by some previous theories as the consequence of variance compensation (48, 49). We demonstrate that the particular form of variance compensation assumed in BLO, when applied to the truncated log-odds, can come close to minimizing the deviation between objective and subjective probabilities (Supplement S12).

In the analyses below we will test whether any or all of these three assumptions of BLO are needed to describe the probability distortion in human behavior. It is easy, for example, to imagine a variant of BLO without variance compensation. However, human performance indicates that something like variance compensation is needed to account for data.

### Overview of the experimental tests of BLO

To test BLO, we first performed a new experiment where each participant completed both a decision-making under risk (DMR) task and a judgment of relative frequency (JRF) tasks. We also re-analyzed the data of Gonzalez and Wu’s (14) DMR experiment. Objective probabilities in these two representative tasks can be readily manipulated and subjective probabilities precisely estimated.

In Gonzalez and Wu (14), 10 participants were tested on 165 two-outcome lotteries, a factorial combination of 15 value sets by 11 probabilities (see Methods). Participants chose between lotteries and sure rewards so that their *certainty equivalent* (CE)—the value of sure reward that is equally preferred—to each lottery was measured. We refer to Gonzalez and Wu’s (14) dataset as GW99, the set of lotteries included in which is large and rich enough to allow for reliable modeling on the individual level—as demonstrated in Gonzalez and Wu (14).

We refer to our new experiment as Experiment JD (see Methods). In the experiment, each of 75 participants completed a DMR task whose procedure and design (Figure S1a) followed that of Gonzalez and Wu (1999) as well as a JRF task (Figure S1b) where participants reported the relative frequency of black or white dots among an array of black and white dots. The same 11 probabilities were used in the two tasks. By comparing the performance of individuals in two different tasks that involved the same set of probabilities, we hoped to identify the possible common representation of probability and how it may vary with task.

Based on the measured CEs (for DMR) or estimated relative frequencies (for JRF), we performed a non-parametric estimate and model fits for the probability distortion of each participant and each task (see Methods and Supplements S4 & S5). Similar to previous studies of DMR (14) and JRF (24, 50), we found inverted-*S*-shaped probability distortions for most participants but also marked individual differences in both tasks (Figure 2abc). About 10% of participants had *S*-shaped (not inverted *S*-shaped) probability distortions. The DMR results of GW99 (Figure 2a) and Experiment JD (Figure 2b) were similar and were collapsed in further analysis whenever possible.

**Figure 2.**
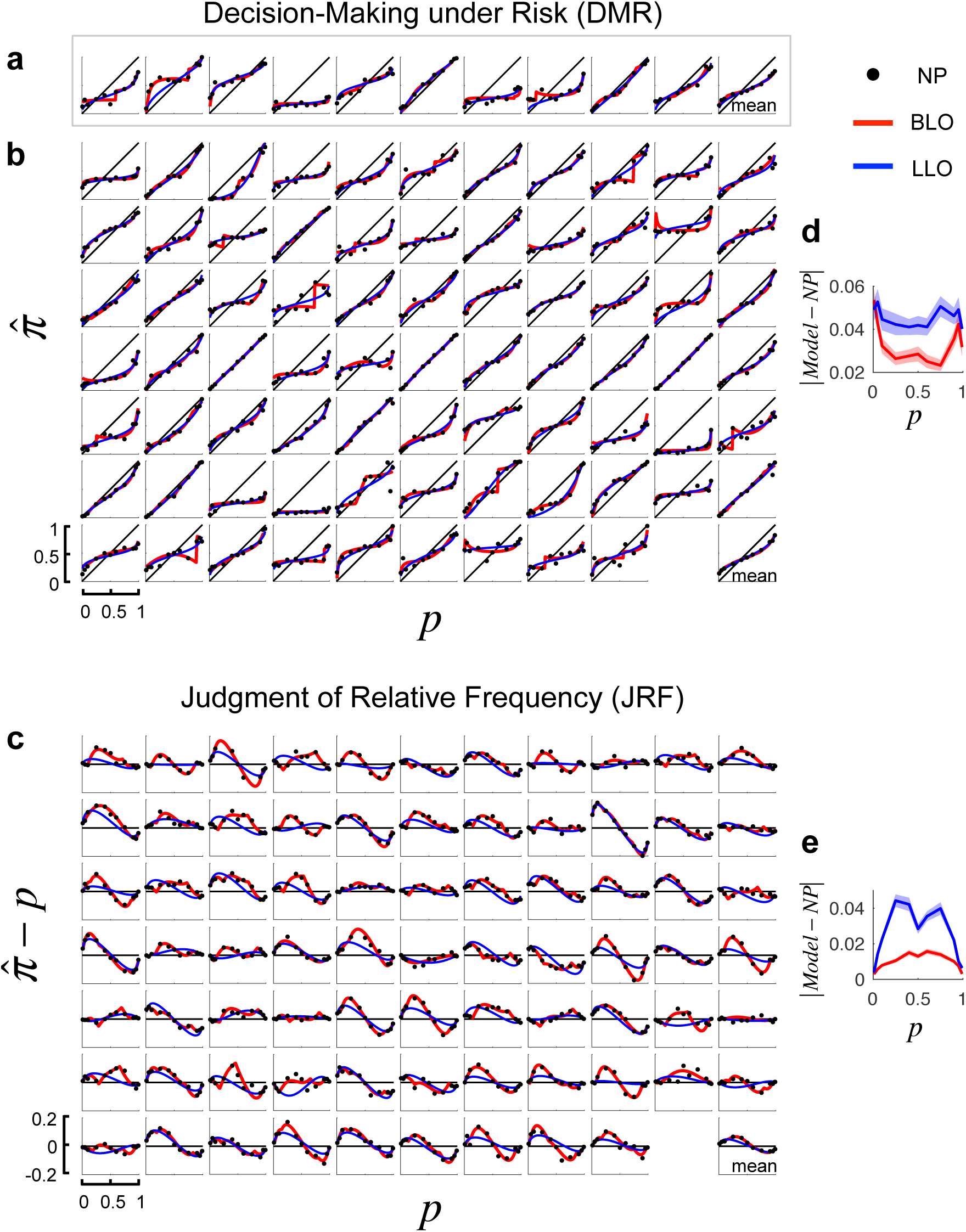
Comparison of model fits to non-parametric estimates of probability distortions. a. Reanalysis of DMR data from Wu & Gonzalez (1999). In the first 10 panels the non-parametric (NP) estimates 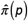 for each participant is plotted versus *p* as black circles. The LLO fit to the participant’s data is drawn as a blue contour, the BLO fit as a red contour. The last panel is the mean across participants. **b. DMR data from our experiment**. The format is identical, with non-parametric estimates and model fits for 75 participants. The last panel is the mean across participants. To make the parametric and non-parametric estimates of probability distortion function comparable, the BLO and LLO fits presented here used the same utility function estimated from the non-parametric estimates of probability distortion. **c. JRF data from our experiment**. For each of the 75 participants we plot the residuals 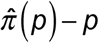 versus *p* to illustrate the small but patterned probability distortions found. We also plot the fits of LLO (blue) and BLO (red) to the residuals. Corresponding panels in **b** and **c** are for the same participant. Compared to the LLO fits (blue curves), the BLO fits (red curves) were overall in better agreement with the non-parametric estimates of probability distortions. **d. e**. Mean absolute deviations of the model fits from the non-parametric estimates are plotted against. *p*, separately for DMR (**d**) and JRF (**e**). Shadings denote SE

We used the non-parametric estimates to assess participants’ probability distortion function and compared model fits with the non-parametric estimates. For an average participant (the last panels in Figure 2abc), the LLO and BLO models provided almost equally good fits. However, an examination of individual participants’ probability distortion revealed that, compared to the LLO fit, the BLO fit captured observed individual differences considerably better. This observation can be quantified using the mean absolute deviations of the model fits from the non-parametric estimates (Figure 2de), which was significantly smaller in BLO than in LLO for 8 out of 11 *p*’s of DMR (paired *t*-tests, *P* < 0.044) and for 10 out of 11 *p*’s of JRF (paired *t*-tests, *P* < 0.005).

### Factorial model comparison

BLO is built on three assumptions: log-odds representation, boundedness, and variance compensation. To test these assumptions, we used *factorial model comparison* (36) and constructed 12 models whose assumptions differ in the following three “dimensions” (see Supplement S6 for details).

#### D1: scale of transformation

The scale of transformation can be the log-odds scale, the Prelec scale (51), or the linear scale based on the *neo-additive family* (52-54, see 55 for a review).

#### D2: bounded versus bounds-free

#### D3: variance compensation

The variance to be compensated can be the encoding variance that varies with *p* (denoted *V* (*p*)) or constant (denoted *V* = *const*).

The models we considered are not all nested nor does factorial model comparison (36) require nested models. Both BLO and LLO are special cases of the 12 models, respectively corresponding to [log-odds, bounded, *V* (*p*)] and [log-odds, bounds-free, *V* = *const*].

For each participant, we fit each of the 12 models to the participant’s CEs (for DMR) or estimated relative frequencies (for JRF) using maximum likelihood estimation (see Supplement S4 for details). The Akaike information criterion with a correction for sample sizes, AICc (56, 57), was used for model selection. For a specific model, the ΔAICc was computed for each participant and each task as the difference of AICc between the model and the minimum AICc among the 12 models. Lower ΔAICc indicates better fit.

For both DMR and JRF, BLO was the model of the lowest summed ΔAICc across participants (Figure 3ab). The results were similar for participants in different experiments (Figure S3). To see how well each of BLO’s assumptions behaves compared to its alternatives, we divided the 12 models into model families by their assumptions on D1, D2, or D3 (e.g. the bounded family and the bounds-free family). We first calculated for each model the number of participants best fit by the model (lowest Δ AICc) and the exceedance probability from the group-level Bayesian model selection (58), which is an omnibus measure of the probability that the model is the best model among the 12 models. The summed number of best-fit participants is then plotted for each model family in Figure 3cd. For both DMR and JRF, the assumptions of BLO outperformed the alternative assumptions on each of the three dimensions, with the summed exceedance probability approaching 1.

**Figure 3.**
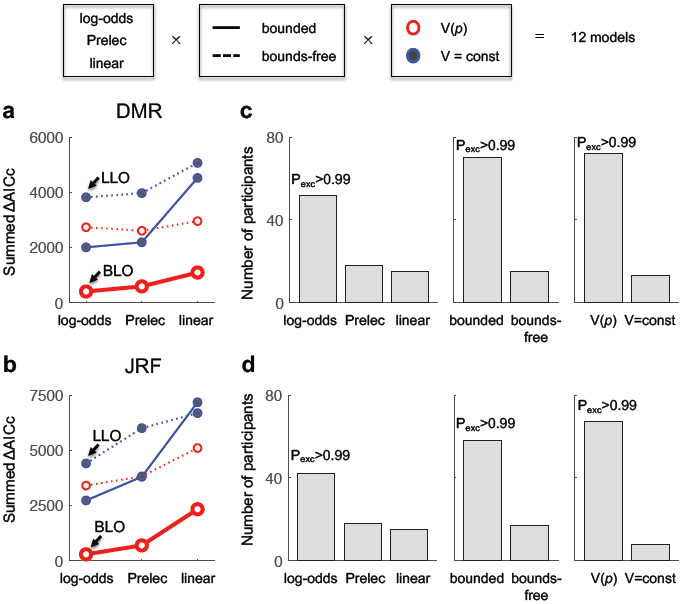
Results of factorial model comparison. We compared 12 models that differ on three dimensions (“factors”) of assumptions: scale of transformation (log-odds, Prelec, or linear), boundedness (bounded or bounds-free) and variance compensation (*V* (*p*) or *V* = *const*). BLO corresponds to [log-odds, bounded, *V* (*p*)]. LLO corresponds to [log-odds, bounds-free, *V* = *const*]. The summed ΔAICc across participants is plotted for each model, separately for DMR (**a**. 85 participants) and JRF (**b**. 75 participants). Lower values of ΔAICc are better. BLO outperformed the alternative models in both tasks. **c, d**. Each assumption of BLO (log-odds, bounded, and *V* (*p*)) also outperformed the alternative assumptions on its dimension. Each panel is for comparisons across one dimension, separately for DMR (**c**) and JRF (**d**). For a family of models with a specific assumption, shaded bars denote the number of participants best accounted by the model family. The P_exc_ above the highest bar denotes the summed exceedance probability of the corresponding model family.

We also performed model comparisons separately for participants with inverted *S*-shaped and participants with *S*-shaped distortions (Figure S4), tested a range of additional models of decision under risk outside the framework we currently used (Figure S5), and tested additional models and an additional dataset (Experiment 1 of Ref. 24) for JRF (Figures S9 & S10). Again, the BLO model outperformed all alternative models (see Supplements S7 & S8).

### Thurstone capacity as a personal signature

According to BLO, the amount of information that the bounded Thurstone scale can encode at a time is limited by 2Ψ /*σ* _Ψ_, where Ψ, as before, is the half-range of the Thurstone scale, and *σ* _Ψ_ denotes the standard deviation of the Gaussian noise on the Thurstone scale. We call Ψ *σ* _Ψ_ the Thurstone capacity. Is the same individual’s Thurstone capacity invariant across tasks?

The Ψ of a specific participant was estimated as a free parameter of BLO from the participant’s reported relative frequency (JRF) or CE (DMR). The value of *σ* _Ψ_ was not fully accessible (see Supplement S11) and we used the estimated BLO parameters, *σ*_*λ*_ (JRF) and *σ* _*CE*_ (DMR), as its surrogates, which respectively characterize the noise variability in the subjective log-odds of JRF and in the CE of DMR. Invariance of Ψ *σ /*_Ψ_ should imply a positive correlation between a participant’s Ψ /*σ*_*λ*_ in JRF and the participant’s Ψ *σ /*_*CE*_ in DMR. In Experiment JD where 75 participants were tested on both tasks, such positive correlation was indeed found (Figure 4), Spearman’s *r*_*s*_ = 0.40, right-tailed *P* < 0.001.

**Figure 4.**
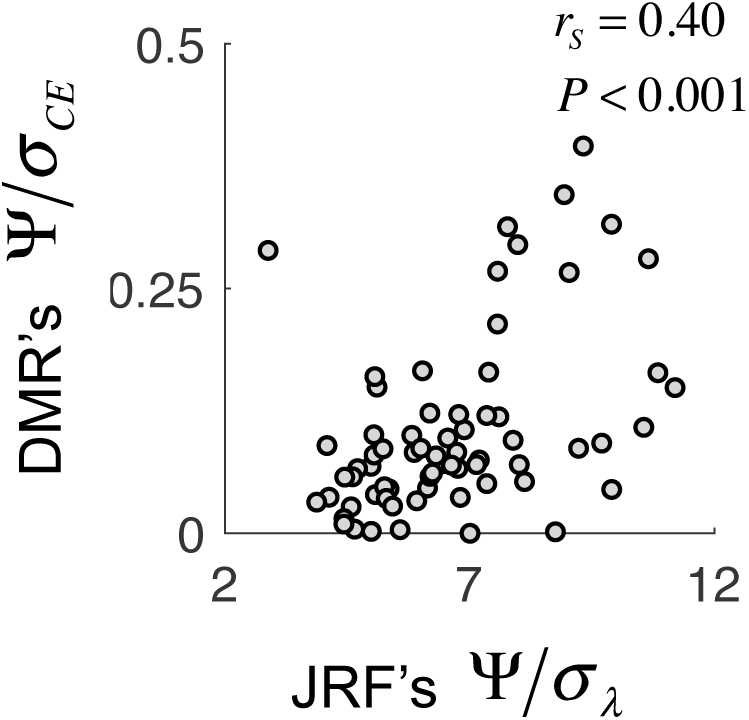
Thurstone capacity as a personal signature. The Ψ /*σ*_*λ*_ estimated in JRF was positively correlated with the Ψ /*σ* _*CE*_ estimated in DMR, for participants who completed both tasks. Each circle is for one participant. (7/75 data points are outside the plot range.) The *r*_*s*_ on the plot refers to Spearman’s correlation coefficient, which is robust to outliers, and *P* is right-tailed.

Given that the two tasks involve entirely different responses and processing of probability information, the across-task correlation between Ψ /*σ*_*λ*_ and Ψ /*σ* _*CE*_ is surprising. In fact, except for modest correlations for Ψ (*r*_*s*_ = 0.23, right-tailed *P* = 0.026) and for the crossover point *p*0 in LLO (*r*_*s*_ = 0.23, right-tailed *P* = 0.025), no positive correlations were found between the two tasks for any other parameters of probability distortion derived from BLO or LLO (see Table S3).

In Experiment JD, 51 participants completed two sessions on two different days, for whom we could also evaluate the correlations of probability distortion parameters across time. Positive correlations were found between Session 1’s and Session 2’s Ψ/ *σ*_*λ /CE*_ (Figure S6) in both the DMR (*r*_*s*_ = 0.60, right-tailed *P* < 0.001) and JRF (*r*_*s*_ = 0.56, right-tailed *P* < 0.001) tasks. Positive across-session correlations for Ψ were also found in both tasks (Figure S6, DMR: *r*_*s*_ = 0.57, right-tailed *P* < 0.001; JRF: *r*_*s*_ = 0.83, right-tailed *P* < 0.001). The Ψ /*σ*_*λ*_ */CE* and Ψ were the only two measures whose across-task and across-session correlations were all significantly positive, among a total of 12 measures derived from BLO or LLO (see Table S3).

These correlations suggest that the Thurstone capacity defined in BLO can be a personal signature that constrains the individual’s probability distortion functions across time and tasks. Meanwhile, the lack of direct access to *σ* _Ψ_ did not allow us to conclude whether Ψ /*σ* _Ψ_ is invariant or only correlated across tasks, which still awaits future empirical tests.

### Maximizing mutual information

The limited Thurstone capacity imposes a trade-off: The wider the interval [Δ^−^, Δ^+^] to encode, the larger the random noise on the encoded values (Figure 1b). In all the datasets we tested, the [Δ^−^, Δ^+^] estimated from participants’ behavior corresponds to a probability range far narrower than the range of objective probabilities ([0.01,0.99]). As we will see below, participants’ choice of [Δ^−^, Δ^+^] maximizes the mutual Shannon information between objective probabilities and their internal representations, a form of efficient encoding (44).

The efficiency of encoding can be quantified by the mutual information between stimuli *s*_1_,…,*s*_*n*_ and responses *r*_1_,…,*r*_*n*_:

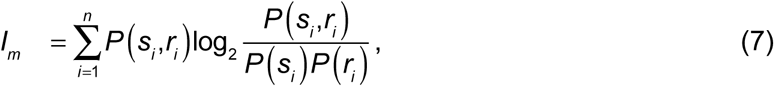

where *P* (*s*_*i*_) denotes the probability of occurrence of a specific stimulus *s*_*i*_, *P* (*r*_*i*_) denotes the probability of occurrence of a specific response *r*_*i*_, and *P* (*s*_*i*_, *r*_*i*_) denotes the conjoint probability of the co-occurrence of the two. Stimuli and responses refer to objective and subjective relative frequencies or probabilities. For a specific task and BLO parameters, we used the BLO model to generate simulated responses and then computed expected mutual information using a Monte Carlo method (see Supplement S10).

For a virtual participant endowed with median parameters, we evaluated how the expected mutual information in JRF or DMR varied with Δ^−^ and Δ^+^, the other parameters being the same. We found that the expected mutual information varied non-monotonically with the values of Δ^−^ and Δ^+^ (Figure 5ab) and for both DMR and JRF, the observed median values of Δ^−^ and Δ^+^ (marked by red circles) were close to the values maximizing the expected mutual information: the mutual information associated with the observed Δ^−^ and Δ^+^ were lower than maximum only by 1.31% for JRF and by 2.83% for DMR. In contrast, if no bounds had been imposed on the probability range of [0.01,0.99] (i.e. Δ^−^ = −4.6, Δ^+^ = 4.6), the mutual information would be 16.7% and 12.3% lower than maximum, respectively for JRF and DMR.

**Figure 5.**
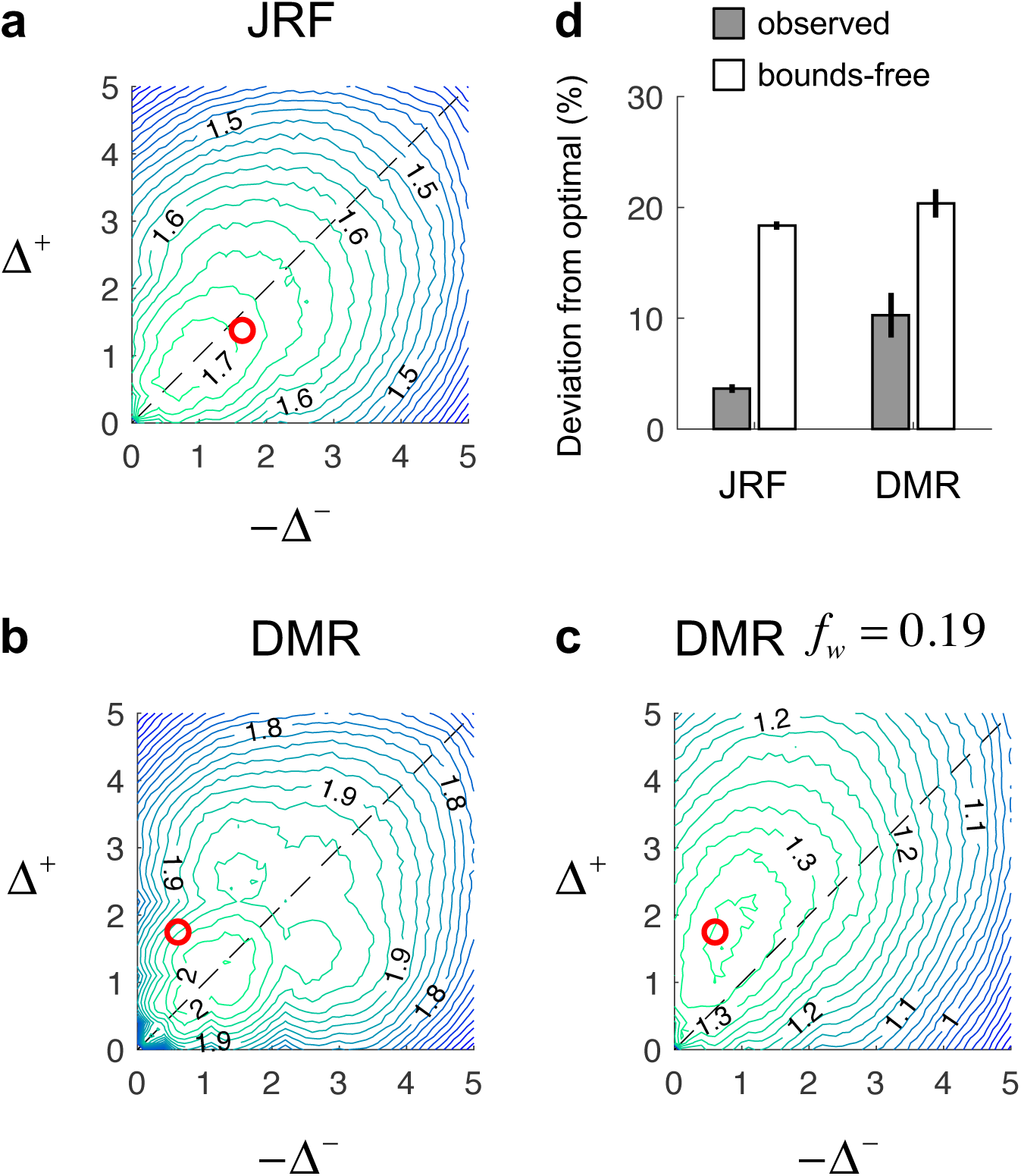
Choice of bounds parameters Δ^−^ and Δ^+^ as mutual information maximization. **a**. Expected mutual information between objective and subjective probabilities (in bits) is plotted against −Δ^−^ and Δ^+^ as contour map for JRF. Higher values are coded as more greenish and lower values as more bluish. **b**. Expected mutual information contour map for DMR. For both tasks, the observed median (−Δ^−^, Δ^+^) (marked by the red circle) was close to maximizing expected mutual information. **c**. When additional multiplicative noise was assumed for the internal representation of probability, the observed asymmetry of Δ^−^ and Δ^+^ in DMR can be explained by maximizing mutual information. The SD of the multiplicative noise was assumed to be 0.19 times of the internal representation of probability. **d**. Individual participants’ percentage of deviation from optimal in expected mutual information for observed (Δ^−^,Δ^+^)versus bounds-free representations. Bars denote mean percentage across participants. Error bars denote SE.

The observed Δ^−^ and Δ^+^ in JRF (Figure S7b) were almost symmetric around 0 (median –1.64 and 1.38, see red circle in Figure 5a), though the difference between Δ^+^ and −Δ^−^ reached statistical significance (Wilcoxon rank sum test, *Z* = –2.21, *P* = 0.027). The observed Δ^−^ and Δ^+^ of the same group of participants in DMR (Figure S7a), however, were highly asymmetric (median –0.60 and 1.75, see red circle in Figure 5b, Wilcoxon rank sum test, *Z* = 3.88, *P* < 0.001), implying that the allocation of representation space in DMR was biased towards larger probabilities. What can explain this asymmetry?

We conjectured that it may also be a consequence of efficient coding, if we take into account the potentially larger noises associated with representing larger expected utilities, (59). That is, a more precise representation is needed for larger probabilities in order to have larger and smaller expected utilities equally discriminable. Indeed, when additional multiplicative noise was assumed for the internal representation of probability (see Supplement S10), we found that the optimal Δ^−^ and Δ^+^ would exhibit the observed asymmetry in DMR (Figure 5c).

We also computed expected mutual information based on individual participants’ BLO parameters and compared each participant’s Δ^−^ and Δ^+^ with the optimal choice suggested by the participant’s Thurstone capacity. Individual participants’ deviation from optimality (Figure 5d) was on average larger than that of the median participant. But still, the observed Δ^−^ and Δ^+^ was only approximately 10% lower than optimality in expected mutual information and much closer to optimality than alternative bounds-free representations.

The results of factorial model comparison reported earlier provided evidence that participants used bounded instead of bounds-free representations. In the mutual information analysis above, we further revealed the rationality behind this boundedness. We found that under their constraint in Thurstone capacity, participants’ choice of the interval to encode was close to maximizing the information transmitted by the Thurstone scale.

### Minimizing expected error

Efficient encoding maximizes the discriminability between subjective probabilities, but cannot guarantee whether the subjective probability is an accurate estimation of the objective probability. For example, suppose the probabilities of hazard for two actions are 0.9 and 0.95 but are estimated to be 0.01 and 0.2, respectively. Though these two actions are well discriminated from each other, decision-making based on such inaccurate subjective estimates can be disastrous.

Polanía et al. (29) assumed that Bayesian decoding follows efficient encoding of value. Similarly, the choice of bounds parameters in BLO only determines how efficiently the truncated log-odds encoded by the Thurstone scale transmits information about the objective probability. The accuracy of the subjective estimate, instead, relies on variance compensation, whose performance is controlled by two parameters of BLO: Λ_0_ and *κ*. The final estimate of log-odds is a weighted average of the truncated log-odds 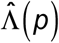 and an anchor Λ_0_ (Eq. 6). The parameter *κ* controls the extent to which the encoding uncertainty influences the weight *ω* _*p*_ for 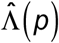 (see Methods). How well did participants choose their variance compensation parameters to improve the accuracy of subjective probabilities?

We define the expected error of subjective estimates as the square root of the mean squared deviations between objective and subjective probabilities for a specific distribution of objective probabilities. Similar to our computation of expected mutual information, we evaluated how the expected error in DMR or JRF varied with Λ_0_ and *κ*, the other parameters being the same (see Supplement S10). We found that the observed Λ_0_ and *κ* for a median participant were close to minimizing expected error, deviating from optimality only by 5.95% for JRF and by 7.67% for DMR (Figure 6ab). The deviation for individual participants was larger (approximately 10% for JRF and 20% for DMR), but still much smaller than representations assuming no variance compensation (Figure 5c).

**Figure 6.**
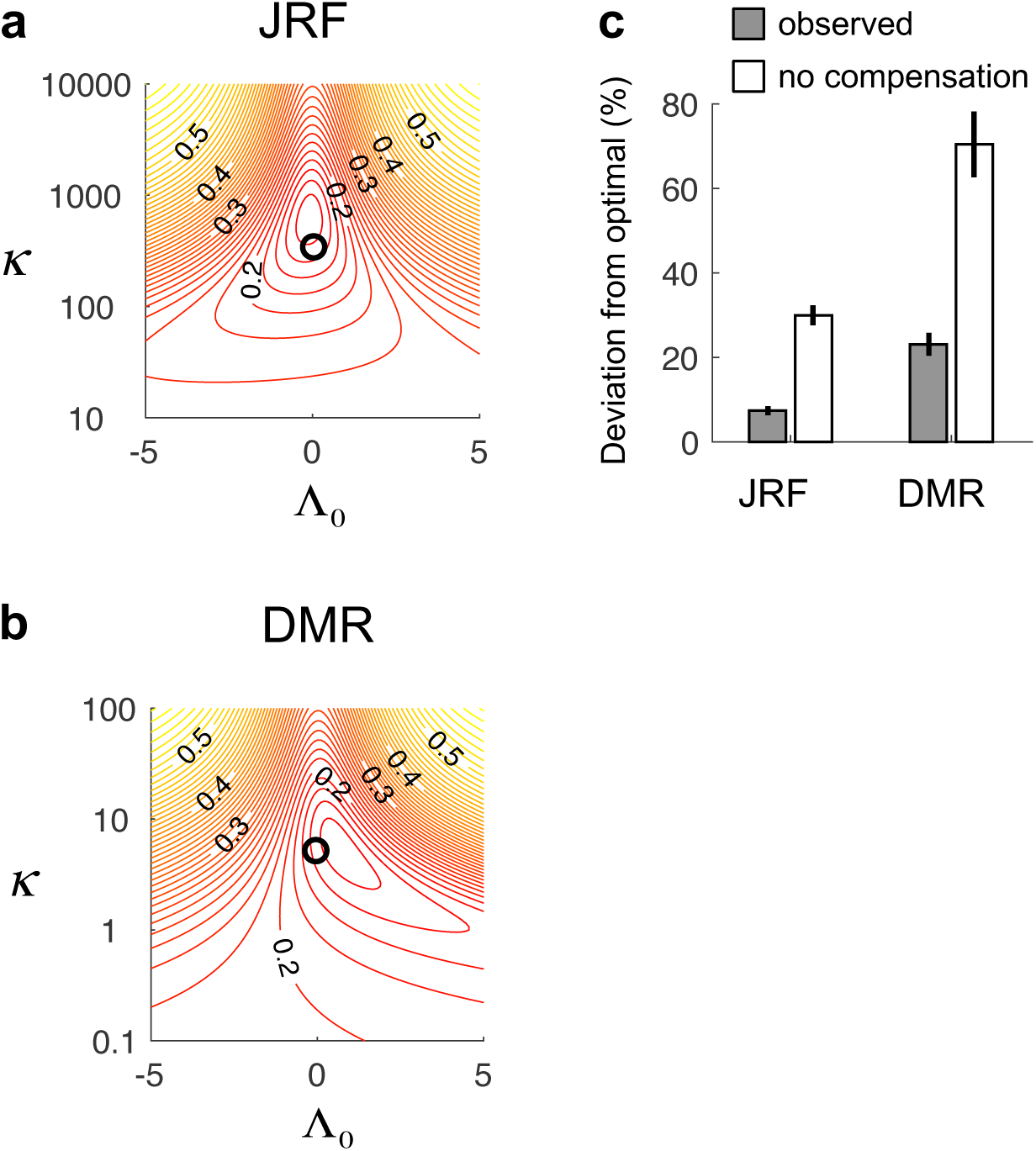
Choice of variance compensation parameters Λ_0_ and *κ* as expected error minimization. **a**. Expected error in probability is plotted against Λ_0_ and *κ* as contour map for JRF. Smaller errors are coded as more reddish and larger errors as more yellowish. **b**. Expected error contour map for DMR. For both tasks, the observed median (Λ_0_, *κ*) (marked by the black circle) was close to minimizing the expected error in probability. **c**. Individual participants’ percentage of deviation from optimality in expected error for observed (Λ_0_, *κ*) versus representations assuming no variance compensation. Bars denote mean percentage across participants. Error bars denote SE.

Finally, we caution that the close-to-optimal choices of parameters we identified above did not necessarily imply neural computations of optimal solutions. They could just follow some simple rules. For example, participants’ choice of Λ_0_ in both tasks was close to 0. In DMR, this choice was actually closer to the mean of the objective log-odds than to the value of Λ_0_ that minimizes expected error (Figure 5b).

## Discussion

The BLO model is intended to model performance in both DMR and JRF tasks and it is the first model that attempts to do so. It is based on three assumptions: log-odds representation, encoding on a bounded Thurstone scale, and variance compensation. We tested each of these assumptions using factorial model comparison to verify that they are all essential to best predict human behavior. That is, if we replace any assumption by the alternatives we considered, the resulting model is strictly inferior to BLO.

We then compared BLO with all of the other models in the literature intended to account for probability distortion. BLO outperformed all these models in accounting for our experimental results as well as the data of Gonzalez and Wu (14). Among the models considered, BLO is the best available *descriptive* model of human use of probability and relative frequency information.

We then considered whether BLO is *normative* in a specific sense. We tested whether participants chose probability distortions that come close to maximizing the **mutual information** between objective probabilities and their imperfect subjective estimates. Two recent articles use the same criterion (maximum mutual information) to model human encoding of value (29) or to re-interpret the context effects of decision under risk (46). These articles taken together are consistent with a claim, supported by considerable experimental data, that many observed failures in DMR can be viewed as attempts to compensate for immutable limits in cognitive processing in order to preserve Shannon information, a form of bounded rationality (37).

### A single model for probability distortion

There are many theoretical models intended to account for inverted-*S*- or *S*-shaped probability distortion: the power model of proportion judgment (60, 61), the support theory model of probability judgment (48, 62), the calibration model (63), the stochastic model of confidence rating (38, 39), and the adaptive probability theory model of decision under risk (49). However, almost all these models were proposed for one specific type of task and not intended as general explanations for observed distortion of probability and relative frequency. Neither do they explain why participants exhibit different probability distortions in different tasks or task conditions. There was even a belief, at least in decision under risk, that the parameters of distortion should be specific to each participant but constant across all tasks (64).

In contrast, BLO models a common mechanism underlying all probability distortion, where we identified one constraint—limited information processing capacity (the Thurstone capacity)—that is pervasive in models of cognitive and perceptual tasks (36, 45, 65) and that may be invariant across different tasks. The probability distortion functions are determined by the constraint as well as close-to-optimal choices under this constraint. We next describe some of the implications of BLO.

### Discontinuities at p = 0 and p = 1

BLO and any model based on the boundedness assumption predict that *π* (0) > 0 and *π* (1) < 1, that is, probability distortion with discontinuities at *p* = 0 and *p* = 1. Such discontinuities are also found in the neo-additive family of weighting functions (55), but are not found in other, widely accepted families of probability distortion such as LLO (14, 24) and Prelec’s family (51). Kahneman and Tversky’s original Prospect theory (Figure 4 of Ref. 12) included similar discontinuities in probability weighting functions.

The bounded ranges of probability represented on the Thurstone scale according to the BLO model fits are fairly limited, approximately [0.16, 0.80] in JRF and [0.35, 0.85] in DMR. Given that the occurrence of probabilities as extreme as 0.05 and 0.95—or even 0.01 and 0.99—is not uncommon in laboratory tasks or real life, bounding is likely to exert detectable influences on probability representation and performance under many circumstances. Indeed, there are clues indicating boundedness in previous studies. For example, Yang and Shadlen (66) studied monkeys’ probabilistic inference and found that the strength of a specific evidence perceived by the monkey was, in general, proportional to the objective log-odds of the evidence. But for “sure evidence” that corresponded to minus or plus infinity in log-odds, the subjective log-odds were bounded, equivalent to [0.15, 0.81] and [0.30, 0.64] in probability for the two tested monkeys.

### Compensation for encoding uncertainty

Anchoring—as a way to explain the inverted-*S*-shaped curve and its individual differences— has been assumed in a few theories or models of probability distortion (48, 49, 67). It can be a way to improve the accuracy of probability judgment, following the perspective of Bayesian inference (47). What distinguishes BLO from previous models is the assumption that anchoring implements compensation for encoding uncertainty. Intuitively, percepts of lower uncertainty should be less discounted and those of higher uncertainty more discounted. If the uncertainty of a percept varies with the value of probability it encodes, so will the reliability weight endowed to the percept.

For the JRF task, the uncertainty may arise from a sampling process, analogous to the sampling in perceptual tasks such as motion perception (68, 69) and pattern detection (70). For the DMR task, where probability is explicitly defined and no explicit sampling process seems to be involved, we still found that the slope of probability distortion relies on a *p* (1− *p*) term, varying with *p* (see Methods). It is as if people are compensating for the variation of a virtual sampling process (49, 71, 72), or for the variation caused by Gaussian noise on the Thurstonian log-odds scale (Supplement S1, see also 73). Lebreton et al. (73) show that a generalized form of *p* (1− *p*) is correlated with the confidence of value or probability perception and is automatically encoded in the ventromedial prefrontal cortex (vmPFC) of the human brain. Under certain circumstances, such variance compensation may result in counterintuitive non-monotonic probability distortion that is indeed empirically observed (see Supplement S13).

### Predicting the slope of probability distortion

Mutual information maximization requires the encoded interval to scale with the range of probabilities in the stimuli. When a narrower interval is encoded, the truncated log-odds encoded on the bounded Thurstone scale for the same objective probability can be more extreme, leading to probability distortion of a steeper slope. Thus BLO predicts that the narrower the probability range of the stimuli, the steeper the slope of distortion.

We performed the following meta-analysis on previous DMR studies to test this prediction. Fox and Poldrack (74, Table A.3) summarized the results of a number of decision-making studies that were modeled in the framework of Prospect Theory. In Fox and Poldrack’s list, we identified the studies where the gamble set was explicitly defined and each gamble consisted of two outcomes that could be denoted (*x*_1_,*p*;*x*_2_,1− *p*) (see Supplemental Table S4 for the 12 studies included). Though different functional forms—LLO, one-parameter and two-parameter Prelec functions (51), and Tversky and Kahneman’s weighting function (13)—had been assumed in different studies, all had a parameter for the slope of probability distortion that is roughly equivalent to the *γ* in LLO. For each study, we computed the standard deviation of objective log-odds and found that, consistent with the BLO prediction, this measure was negatively correlated with the slope of probability distortion (Figure 7a), *r*_*s*_ = –0.56, left-tailed *P* = 0.030. Assuming optimal choice of Δ^−^ and Δ^+^, we further quantitatively predicted the slope of distortion for each study, which resembled the observed slopes (Figure 7b, see Supplement S14 for details).

**Figure 7.**
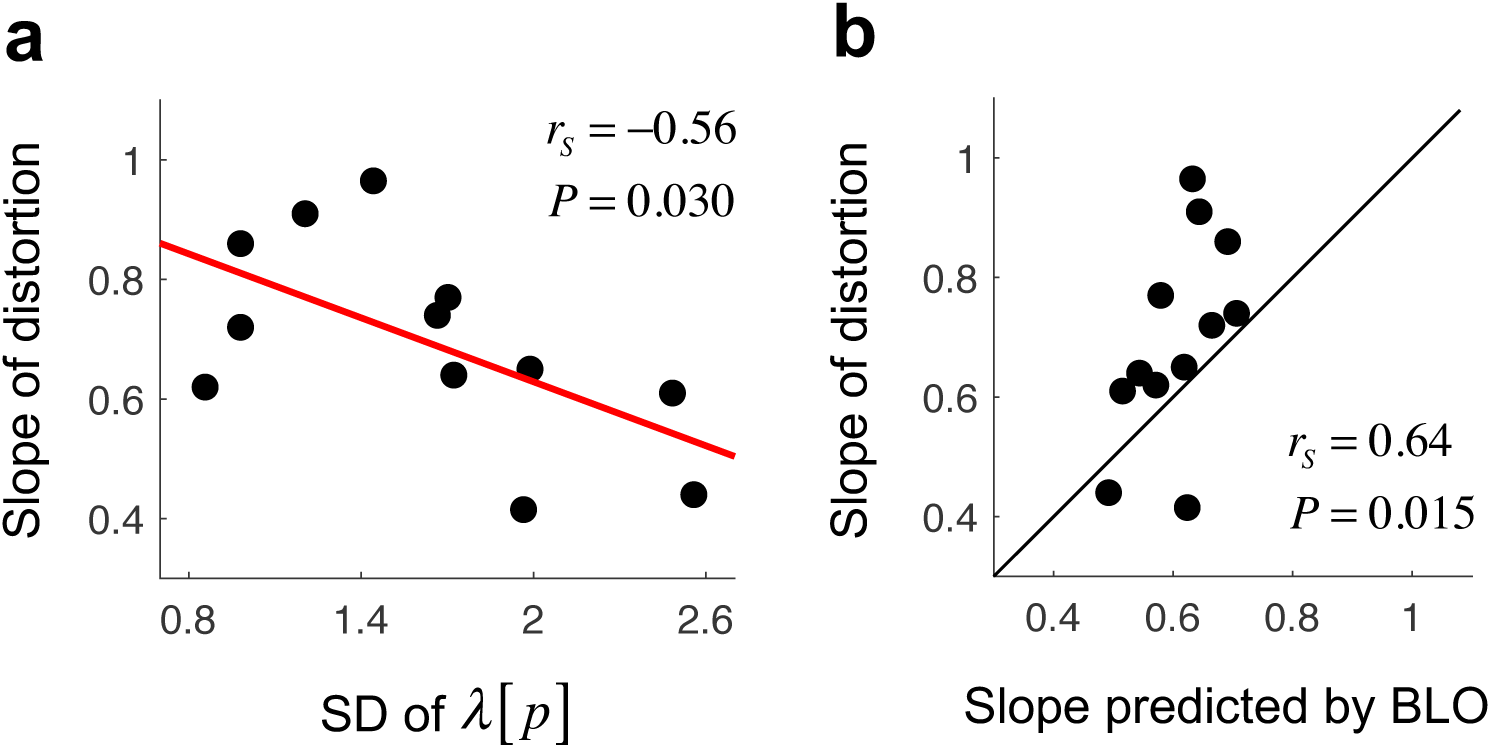
Meta-analysis of previous studies supporting BLO’s prediction on the slope of probability distortion in decision under risk. **a**. The estimated slope of probability distortion 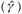 is plotted against the standard deviation of the objective log-odds (*λ* (*p*)) of the gamble set, where *p* denotes the probability for the higher outcome of a two-outcome gamble, (*x*_1_,*p*;*x*_2_,1− *p*). Each data point is for one published study. The red line denotes the regression line. The correlation is negative and significant. We describe the selection of studies in the text. See Table S4 for a full list of the studies. That the slope of distortion decreases with the standard deviation of *λ* (*p*) is consistent with the prediction of BLO. **b**. Estimated slope of probability distortion is plotted against the slope predicted by BLO for each study (see Supplement S14 for details).

### The crossover point

A puzzle we did not fully address earlier concerns the crossover point of probability distortion (i.e. the point on the distortion curve where overestimation changes into underestimation or the reverse). It has been frequently observed that the crossover point is near 0.5 for the JRF task (24) but approximately 0.37 for the DMR task (51). That is, the probability distortion is symmetric around 0.5 in the former but asymmetric in the latter. There are plausible reasons to have symmetry, but why asymmetry? Here we conjecture that the asymmetry is also driven by the maximization of mutual information, which, for the DMR task, relates to having the CEs of different gambles as discriminable as possible. Following conventions (14, 74) and for parsimony, we had assumed a uniform Gaussian noise on the CE scale. However, larger CEs may tend to be associated with higher variances, an analogue to Weber’s law (59). To compensate for this, more of the representational scale should be devoted to larger probabilities and thus to the larger CEs associated with them. Indeed, we found that the less-than-0.5 crossover point in DMR is associated with bounds [Δ^−^, Δ^+^] that biases towards larger probabilities (see our discussion of asymmetric [Δ^−^, Δ^+^] in Results), which effectively implements such a strategy of probability representation.

### Open questions and future directions

The judgment of relative frequency and decision under risk are the only two tasks where BLO and its assumptions have been tested, but these two tasks together represent a vast body of previous research. The model may be applied to a wider range of tasks involving frequency and probability. Whether it succeeds or fails, it will likely shed light on the common and distinctive mechanisms of probability distortion in different tasks.

What determine the slope and crossover point of probability distortion in a specific task or task condition? Why may the parameters of probability distortion change from task to task and from individual to individual? In the present study we have provided a tentative answer: they change because the brain actively compensates for its own fixed limitations.

We propose probability distortion as a consequence of bounded rationality but must caution that the optimality found on the group level cannot guarantee optimality for every individual. For example, for the anchor parameter of BLO whose optimal value is determined by the prior distribution of probabilities, there were still considerable individual differences. One possibility is that some individuals may be slow to update their prior or even not able to correctly learn the true prior (75). Besides, there are large individual differences in the optimality of using cognitive resources (76).

Important questions for future research also include: How may probability distortion change from trial to trial? We conjecture that the human representation of probability can adapt to the environment, in the spirit of efficient coding (42-44) or Bayesian inference (47). The current version of BLO is a stationary model, whose prediction will not change with time or experience. In contrast, non-stationarity has been identified in probability distortion for both the judgment of relative frequency (24) and decision under risk (77).

We chose not to consider “decision from experience” (20)—another important form of decision-making—because the decision from experience task does not require that the decision maker estimate the frequency of items (19, 78). The decision maker may estimate the multinomial distribution of rewards in a card deck—or she may simply register reward and punishment and base her decision on a form of reward averaging or reinforcement learning. The results of a comprehensive model competition (79) are consistent with this claim. More recently, there has been neuroimaging evidence that human decisions from experience may be based on the retrieval of individual samples from past experience (80, 81). If the decision maker does not estimate relative frequency then BLO does not apply.

A final note: Kahneman and Tversky’s original Prospect Theory contained the assumption that decision makers would first interpret (“edit”) available information (12). In this initial *editing* stage they might, for example, convert the probability 0.31317 to the more tractable 1/3. Only then would they assign prospect values to lotteries in the second, *evaluation* stage. In presenting the BLO model we focus on evaluation. Still, nothing about the theory would preclude adding an editing phase or discretizing the representation of probability if justified by empirical results.

## Methods

### Experiment

Experiment JD was approved by the Institutional Review Board of School of Psychological and Cognitive Sciences at Peking University. All participants gave written informed consent in accordance with the Declaration of Helsinki. Each participant performed two tasks: Decision-Making under Risk (DMR) and Judgment of Relative Frequency (JRF).

The procedures and designs of the DMR task were the same as those of Gonzalez and Wu (14), except that payoffs in the gambles were in RMB instead of in USD. On each trial (Figure S1a), participants were presented with a two-outcome gamble (*x*_1_,*p*;*x*_2_,1− *p*) and tables of sure amounts of rewards. They were asked to check on each row of the tables whether they preferred the gamble or the sure amount. The range of the sure amounts started with [*x*_2_,*x*_1_], and was narrowed down in the second table so that we could estimate participants’ certainty equivalent (CE) for the gamble. There were 15 possible outcome pairs (*x*_1_,*x*_2_): (25, 0), (50, 0), (75, 0), (100, 0), (150, 0), (200, 0), (400, 0), (800, 0), (50, 25), (75, 50), (100, 50), (150, 50), (150, 100), (200, 100), (200, 150). There were 11 possible probabilities: 0.01, 0.05, 0.1, 0.25, 0.4, 0.5, 0.6, 0.75, 0.9, 0.95, 0.99. A full combination of them resulted in 165 different gambles used in the experiment.

The stimuli and procedures of the JRF task followed Zhang and Maloney (24). On each trial (Figure S1b), participants were presented with an array of black and white dots and reported their estimate of the relative-frequency of black or white dots by clicking on a horizontal bar with tick marks from 0 to 100%. Each participant was randomly assigned to report the relative frequency either for the black or for the white dots. The objective relative frequency of JRF was chosen from the same 11 possible values as its counterpart in DMR. The total number of dots (numerosity) in a trial was varied across trials, which could be 200, 300, 400, 500, or 600. The dots in each display were distributed within a circular area of 12° diameter or a square area of 17°×17° diameter.

Experiment JD (a total of 75 participants) consisted of two sub-experiments, JDA (51 participants, 20 male, aged 18 to 29) and JDB (24 participants, 10 male, aged 18 to 27). Six additional participants failed to complete the experiment for technical or personal reasons. Each session had 11 (probability) × 15 (outcome pair) = 165 DMR trials and 11 (probability) × 5 (numerosity) × 6 = 330 JRF trials, which took approximately two hours. In Experiment JDA, each participant completed two sessions on two different days, so that we could evaluate the consistency of their performance. Trials from the two tasks were randomly interleaved. In Experiment JDB, each participant completed only one session, during which one task preceded the other, with DMR first for half of the participants and JRF first for the other half. Similar patterns of probability distortions (Figure 2bc, first 51 panels for Experiment JDA and last 24 panels for Experiment JDB) and results of model comparisons (Figure S3) were found for participants in the two sub-experiments. Thus we collapsed the two sub-experiments in our analysis whenever applicable.

### Applying BLO to JRF

We need additional assumptions when applying BLO to the JRF experiments. One of the key assumptions of BLO is variance compensation and, to apply BLO, we need to specify a model of the participant’s sampling process and the variance of the resulting estimates. First, we assume that humans may not have access to all the tokens presented briefly in a display or in a sequence, due to perceptual and cognitive limits (82, 83). Instead, they take samples from the population and are thus subject to the randomness associated with sampling. Within BLO, probability distortion arises in part from a compensation for the sampling noise captured in our model by the reliability parameter *ω* _*p*_.

Denote the total number of dots in a display as *N* and the relative frequency of black dots as *p*. Suppose a sample of *n*_*s*_ dots is randomly drawn from the display. We assume that the sampling is without replacement (see Supplement S8 for models with the alternative assumption of sampling with replacement). That is, the same dot will not be drawn twice during one sampling, which is reasonable in our case. As a result, the variance of 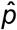 requires a correction for finite population (84, see Supplement S2 for the derivation):

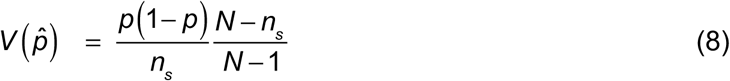

The finite population correction is intuitive: the larger the sample size relative to the population, the smaller the variance. When *n*_*s*_ = *N*, i.e. when the whole population is included in the sample, we should have 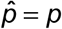 for each sample and thus 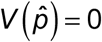. At the other extreme, when *n*_*s*_ = 1, sampling without replacement is equivalent to sampling with replacement, the familiar *p* (1− *p*). For any participant and numerosity condition, when *n*_*s*_ > *N*, that is, when the participant was able to sample all the dots in the display, we forced 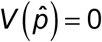.

The BLO variance correction is a weighted mixture of an estimate based on the sample and an “anchor” Λ_0_(Eq. 6), with the weight for the former

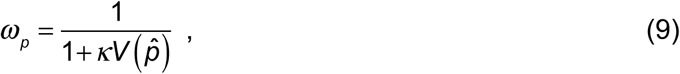

where *κ* > 0 is a free parameter. See Supplement S12 for how this form of *ω*_*p*_ can serve as an approximate solution to minimizing the expected error in estimating probability.

Finally, we modeled participants’ reported relative frequency *π* (*p*) as a function of *p* perturbed by additive Gaussian error:

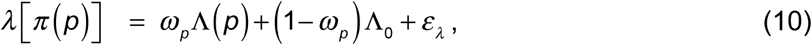

where *ε*_*λ*_ is Gaussian error on the log-odds scale with mean 0 and variance 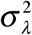.

### Applying BLO to DMR

To model *π* (*p*), BLO’s assumptions for different tasks are the same, except that encoding variance is task-specific. Probability is described explicitly in DMR and there seems to be no uncertainty about its value. Participants’ choices suggested, however, that they were still compensating for some kind of encoding uncertainty that varies with the value of probability. Gaussian encoding noise on the Thurstone scale in log-odds, when transformed back to the probability scale, results in variance that is approximately proportional to *p*(1 − *p*) (see Supplement S1 for proof). The reliability parameter in Eq. 6 is thus:

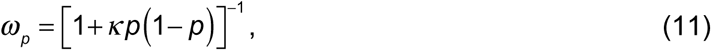

where *κ* > 0 is a free parameter. This same equation can be reached if, alternatively, we assume that participants were compensating for a virtual sampling process (the 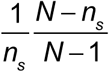 term in Eq. 8 can be assimilated into *κ* for constant *N* and *n*_*s*_). Compensation for virtual sampling was assumed in some previous theories on probability distortion (49, 71, 72).

Any lottery in GW99 or Experiment JD can be written as (*x*_1_,*p*;*x*_2_,1− *p*), which offers the value *x*_1_ with probability *p* and otherwise *x*_2_, with *x*_1_ > *x*_2_ ≥ 0. For each participant, we modeled the certainty equivalent (CE) of each lottery using Cumulative Prospect Theory (CPT) (13) and assumed a Gaussian error term on the CE scale, as in Gonzalez and Wu (14):

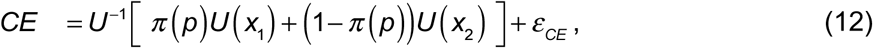

where *U* (·) denotes the utility function, *U* ^−1^ (·) denotes the inverse of *U* (·), *π* (*p*) denotes the probability distortion function (same as that in Eq. 10, except without Gaussian error), and *ε*_*CE*_ is a Gaussian random variable with mean 0 and variance 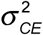. The utility function for non-negative gains alone (none of the lotteries involved losses) was assumed to be a power function with parameter *α* > 0:

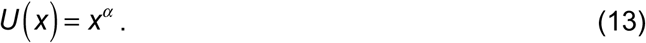

## Acknowledgments

We would like to thank Richard Gonzalez and George Wu for sharing their dataset with us. H.Z. was supported by grants 31571117 and 31871101 from National Natural Science Foundation of China and funding from Peking-Tsinghua Center for Life Sciences. L.T.M. was supported by grant EY019889 from the National Institutes of Health, the Humboldt Research Prize of the Alexander v. Humboldt Foundation, a Guggenheim Fellowship from the John Simon Guggenheim Foundation and a Fellowship from the Institut des Etudes Avancées de Paris. A previous version of this work was published as a preprint on bioRxiv.

## Author contributions

H.Z. and L.T.M. conceived the study. H.Z. and X.R. designed the experiment. X.R. performed the experiment. H.Z. and L.T.M. developed the theory. H.Z. and X.R. analyzed the data. H.Z. and L.T.M. wrote the manuscript.

The authors declare no conflicts of interest.

