## Supplements for "The bounded rationality of probability distortion"

### Supplement S1: Derivation of the encoding variance

Suppose $X$ is an encoding of the probability $p$ on the log-odds scale contaminated by a Gaussian noise: $X \sim Normal\left(\lambda(p), \Sigma\right)$, where $\lambda(p) = \log\frac{p}{1-p}$ and $\Sigma$ is a constant for variance. When transformed back to the probability scale, the representation is $\lambda^{-1}[X]$. Given that $\lambda^{-1}[X] = \frac{1}{1+e^{-X}}$ is a logistic function of the Gaussian random variable $X$, its variance has the following approximation (Daunizeau, 2017):

$$Var\left(\lambda^{-1}[X]\right) \approx \lambda^{-1}\left[\frac{\lambda(p)}{\sqrt{1+\frac{3\Sigma}{\pi^2}}}\right]\left(1-\lambda^{-1}\left[\frac{\lambda(p)}{\sqrt{1+\frac{3\Sigma}{\pi^2}}}\right]\right)\left(1-\frac{1}{\sqrt{1+\frac{3\Sigma}{\pi^2}}}\right). \tag{S1}$$

Let $\zeta = \frac{3\Sigma}{\pi^2}$, the right side of Eq. S1 can be written as a function of $\zeta$:

$$g(\zeta) = \lambda^{-1}\left[\frac{\lambda(p)}{\sqrt{1+\zeta}}\right]\left(1-\lambda^{-1}\left[\frac{\lambda(p)}{\sqrt{1+\zeta}}\right]\right)\left(1-\frac{1}{\sqrt{1+\zeta}}\right). \tag{S2}$$

It is easy to prove that the values of $g(\zeta)$ and its first derivative at $\zeta = 0$ are $g(\zeta)\big|_{\zeta=0} = 0$ and $g'(\zeta)\big|_{\zeta=0} = \frac{1}{2}p(1-p)$. When $\zeta = \frac{3\Sigma}{\pi^2} \ll 1$, the Tayler expansion of $g(\zeta)$ on $\zeta$ is:

$$
\begin{aligned}
g(\zeta) &= g(\zeta)\big|_{\zeta=0} + g'(\zeta)\big|_{\zeta=0}\zeta + o\left(\zeta^2\right) \\
&= \frac{1}{2}p(1-p)\zeta + o\left(\zeta^2\right) \\
&\approx \frac{1}{2}p(1-p)\zeta = \frac{3\Sigma}{2\pi^2}p(1-p)
\end{aligned}
\quad . \tag{S3}
$$

That is, the variance of $\lambda^{-1}[X]$ has the approximate form,

$$
Var\left(\lambda^{-1}[X]\right) \approx \frac{3\Sigma}{2\pi^2}p(1-p). \tag{S4}
$$

See Lebreton, Abitbol, Daunizeau, and Pessiglione (2015) for a similar proof in a different guise.

### Supplement S2: Variance of sampling without replacement

Suppose that $B_1, \cdots, B_{n_s}$ are sampled[1] *without replacement* from a population $\{b_1, \cdots, b_N\}$ of $N \geq n_s$ members taking on only the values 0 or 1. The mean of the population is $p = N^{-1}\sum_{i=1}^{N} b_i$, the proportion of the population that takes on the value 1. The mean of the sample is $P_s = n_s^{-1}\sum_{i=1}^{n_s} B_i$ (a random variable) and the variance of $P_s$ is (Cochran, 1977, Theorem 2.2):

$$\begin{aligned}
&Var(P_s) = E[P_s - p]^2 \\
&= \frac{\sum_{i=1}^{N}(b_i - p)^2}{N-1}\frac{1}{n_s}\frac{N-n_s}{N} \\
&= \frac{\sum_{i=1}^{N} b_i^2 - 2p\sum_{i=1}^{N} b_i + \sum_{i=1}^{N} p^2}{n_s N}\frac{N-n_s}{N-1} \\
&= \frac{Np - 2pNp + Np^2}{n_s N}\frac{N-n_s}{N-1} \\
&= \frac{p(1-p)}{n_s}\frac{N-n_s}{N-1}
\end{aligned} \quad . \tag{S5}$$

The first term $p(1-p)/n_s$ on the right hand side is the variance of samples drawn with replacement. The second term $(N-n_s)/(N-1)$ is the "correction for sampling without replacement". Eq. S5 can be derived alternatively by recognizing that $NP_s$, as a random variable, follows a hypergeometric distribution (Wackerly, Mendenhall, & Scheaffer, 2008).

---

[1] Random variables are denoted by upper case letters, corresponding non-random variables by the same letter in lower case: for example, "the random variable *X* has probability density function *f(x)".*

When $n_s = 1$, sampling with and without replacement are indistinguishable and the sample variance is the familiar

$$Var(B_1) = p(1 - p). \tag{S6}$$

When $n_s = N$, sampling without replacement is exhaustive and the variance is 0 (the sample is always the population and its mean is always precisely the population mean).

### Supplement S3: The relationship between BLO and LLO

We compare LLO (Eq. 2)

$$\lambda\left[\pi(p)\right]=\gamma\lambda(p)+(1-\gamma)\lambda(p_0)$$

to BLO rewritten:

$$\begin{aligned}\Lambda^{\omega}(p)&=\omega_p\Lambda(p)+\left(1-\omega_p\right)\Lambda_0\\&=\omega_p\frac{\Psi}{\left(\Delta^+-\Delta^-\right)/2}\left[\Gamma\left(\lambda[p]\right)-\left(\Delta^-+\Delta^+\right)/2\right]+\left(1-\omega_p\right)\Lambda_0\\&=\tau\omega_p\Gamma\left(\lambda[p]\right)+\left(1-\tau\omega_p\right)\frac{\left(1-\omega_p\right)\Lambda_0-\tau\omega_p\left(\Delta^-+\Delta^+\right)/2}{1-\tau\omega_p}\end{aligned} \quad \text{(S7)}$$

where $\tau\equiv\dfrac{\Psi}{\left(\Delta^+-\Delta^-\right)/2}$. For $\lambda[p]$ within the range $\left[\Delta^-,\Delta^+\right]$, we have: $\Gamma\left(\lambda[p]\right)=\lambda[p]$. Comparing the equations above, over the range $\left[\Delta^-,\Delta^+\right]$ $\tau\omega_p$ replaces $\gamma$ and $\dfrac{\left(1-\omega_p\right)\Lambda_0-\tau\omega_p\left(\Delta^-+\Delta^+\right)/2}{1-\tau\omega_p}$ replaces $\lambda\left(p_0\right)$. In LLO, both $\gamma$ and $p_0$ are fixed but in BLO, however, the reliability parameter $\omega_p$ may vary with the value of $p$ depending on the model of variance appropriate to a given task. If a specific dataset is generated by BLO but fitted by LLO, we would expect that the estimates of $\gamma$ and $p_0$ would change with experimental conditions as predicted by BLO. Consequently, we can fit LLO to data and look for the pattern of deviations in the fitted coefficients predicted by BLO, a test of the BLO Model (e.g. Figure 7).

### Supplement S4: Fitting models to data

For both the JRF and DMR tasks, we fit each model to each individual participant's responses separately using maximum likelihood estimates. We used **fminsearchbnd** (J. D'Errico), a function based on **fminsearch** in MATLAB® (MathWorks) to search for the parameter values that minimized negative log likelihood, subject to limits described below. We estimated the parameter values maximizing likelihood by repeating the search process 1000–2000 times using different, randomly selected starting points and selected the parameter value estimates corresponding to the smallest value in negative log likelihood obtained. For the participants who completed two sessions, we fit the two sessions separately and together, as required by the analyses in the text. See Supplemental Table S1 for a summary of notations.

**Parameter settings for JRF**

1. **Bounded Log-Odds (BLO) Model**. The implementation of the BLO Model in JRF has seven free parameters, including bounds $\Delta^-$ and $\Delta^+$, half-range of the Thurstone scale $\Psi$, variance compensation parameters $\Lambda_0$ and $\kappa$, sample size $n_s$, and noise variance $\sigma_\lambda^2$. Their limits were set as described below.

$\Delta^-, \Delta^+ \in \left[\ \lambda(0.01),\ \lambda(0.99)\ \right]$. Because the relative frequencies of the stimuli were in the range [0.01, 0.99], a $\Delta^-$ less than $\lambda(0.01)$ or a $\Delta^+$ greater than $\lambda(0.99)$ would lead to the same log likelihood as a $\Delta^-$ equal to $\lambda(0.01)$ or a $\Delta^+$ equal to $\lambda(0.99)$.

$\Psi \in (0,\infty)$. In practice, we fit the scaling parameter $\tau \in (0,\infty)$, with $\Psi \equiv \tau\Delta$, where $\Delta \equiv \left(\Delta^+ - \Delta^-\right)/2$.

$\Lambda_0 \in [-10,10]$. The anchoring parameter.

$\kappa \in (0,\infty)$. The parameter that controls the extent to which the encoding uncertainty influences the weight $\omega_p$.

$n_s \in [1, N]$. Although $n_s$ is described as sample size, we treated it as a continuous variable for convenience. Its upper limit was the number of samples in the stimulus. This limit varied from stimulus to stimulus over the range 200 to 600. When the $n_s$ was estimated to be 1, the finite population correction factor in Eq. S5, $\frac{N-n_s}{N-1}$, would equal to 1, thus indistinguishable from a lack of finite population correction.

$\sigma_\lambda^2 \in (0,\infty)$.

2. **Bounded Prelec Model**. Its parameters and the limits on the parameters were the same as BLO, except that the limits for $[\Delta^-, \Delta^+]$ were $[\lambda'(0.01), \lambda'(0.99)]$ where $\lambda'(p) = -\log(-\log p)$, the counterpart of log-odds for Prelec's function.

3. **Bounded Linear Model**. Its parameters and the limits on the parameters were the same as BLO, except that the limits for $[\Delta^-, \Delta^+]$ were [0.01, 0.99], and the limits for $\Lambda_0$ were [0, 1], since all these parameters were now on the probability scale.

In the factorial model comparison reported in Figure 3, we tested a total of 12 models whose parameters were subsets of $\Delta^-$, $\Delta^+$, $\tau$, $\Lambda_0$, $\kappa$, $n_s$, $\sigma_\lambda^2$ and whose limits can be derived from the limits specified above. See main text.

**Parameter settings for DMR**

The parameter settings for the DMR task were the same as those of JRF except that the sampling-related parameter $n_s$ was omitted and there was an additional parameter $\alpha$ for the utility function, whose limits were set to be $(0,\infty)$.

### Supplement S5: Non-parametric estimation of probability distortion

A non-parametric estimation of probability distortion is plotted in Figure 2 for each participant and each task. For JRF, where participants explicitly estimated their subjective relative frequency, the non-parametric estimation $\hat{\pi}_{NP}(p)$ for a specific $p$ was simply the participant's mean estimate across trials, averaged on the log-odds scale:

$$\hat{\pi}_{NP}(p) = \lambda^{-1}\left[\frac{1}{m}\sum_{t=1}^{m}\lambda\left[\hat{\pi}_t(p)\right]\right] \tag{S8}$$

where $\hat{\pi}_t(p)$ denotes the participant's estimate of relative frequency on trial *t*, $t = 1,2,...,m$ .

For the $\hat{\pi}_{NP}(p)$ of DMR, we modeled participants' CE in the framework of CPT as we did for BLO and LLO fits (Eq. 12) except that no functional form was assumed for the probability distortion function. Instead, the $\hat{\pi}_{NP}(p)$ for each of the 11 *p*'s was fitted as a free parameter. The same power functional form was assumed for the utility function in the non-parametric estimation as in the model fits to minimize possible differences irrelevant to probability distortion. This procedure was different from the non-parametric method of Gonzalez and Wu (1999), where no functional forms were assumed for either probability distortion or utility. We verified in the GW99 dataset that our non-parametric estimation of probability distortion led to similar results as Gonzalez and Wu (1999) (Figure S2).

For some participants, when probability distortion was modeled by BLO or LLO, the estimated utility function was very different from assuming the non-parametric probability distortion, because of the joint estimation of the utility and probability distortion functions. For and only for Figure 2, to make the parametric and non-parametric estimations of probability distortions comparable, we performed additional fitting for BLO and LLO where the utility function, using the same participant's exponent $\alpha$ estimated from the non-parametric estimation.

### Supplement S6: Factorial model comparisons

We used *factorial model comparison* (van den Berg, Awh, & Ma, 2014) to separately test the assumptions of BLO, comparing alternative models that differ in the following three "dimensions".

**D1: scale of transformation.** We considered two alternatives to the log-odds scale: the "Prelec scale" and the linear scale.

The Prelec scale is derived from Prelec's function (Prelec, 1998):

$$\pi(p) = \exp\left(-\beta(-\log p)^{\eta}\right) \tag{S9}$$

with $\eta$ and $\beta$ as free parameters. LLO and the Prelec families both are among the probability weighting functions that typically fit best to data (Cavagnaro, Pitt, Gonzalez, & Myung, 2013; Stott, 2006). They are difficult to distinguish empirically (Cavagnaro et al., 2013; Luce, 2000).

Taking logarithms and negating twice on both sides of Prelec's function, we can see that Prelec's function is equivalent to a linear transformation

$$\lambda'\left[\pi(p)\right] = \eta\lambda'[p] - \log\beta \tag{S10}$$

on the Prelec scale

$$\lambda'(p) = -\log(-\log p) \ . \tag{S11}$$

Note that the Prelec scale differs from the log-odds scale in its asymmetry: two probabilities that are symmetric around 0.5 will be symmetric around 0 when transformed to the log-odds scale but will be asymmetric on the Prelec scale.

The functional form of the linear scale is based on the *neo-additive family* (Bell, 1985; Birnbaum & Stegner, 1981; Chateauneuf, Eichberger, & Grant, 2007; Cohen, 1992; Gilboa, 1988; Loomes, Moffatt, & Sugden, 2002; Teitelbaum, 2007; see Wakker, 2010 for a review) which refers to a linear transformation of probability except that it may have discontinuities at the extremes to ensure $\pi(p)$ is within [0,1]:

$$\pi(p)=\begin{cases} 0, & if\ p=0\ or\ kp+b<0 \\ 1, & if\ p=1\ or\ kp+b>1 \\ kp+b, & otherwise \end{cases} \tag{S12}$$

where $k$ and $b$ are free parameters. The linear scale is accordingly defined as

$$\lambda^{\circ}(x)=\begin{cases} 0, & if\ x<0 \\ 1, & if\ x>1 \\ x, & otherwise \end{cases} . \tag{S13}$$

For models that use the Prelec scale, we simply replaced the log-odds and its reverse transformation with $\lambda'(p)$ and its reverse. For models that use the linear scale, the log-odds transformation was replaced by $\lambda^{\circ}(p)$ but there was no inverse transformation, because $\lambda^{\circ}(p)$ is not invertible, and no need for an inverse transformation, because $\lambda^{\circ}(p)$ corresponds to probability.

In BLO, a linear transformation (Eq. 5) was applied to map the bounded interval $\left[\Delta^-,\Delta^+\right]$ to the full range of the Thurstone scale $[-\Psi,\Psi]$, which shifts $\Gamma(\lambda[p])$ by $-\left(\Delta^-+\Delta^+\right)/2$ and then scales it by $\tau\equiv\dfrac{\Psi}{\left(\Delta^+-\Delta^-\right)/2}$. For models that assume bounds-free representations, or for models that assume the Prelec or linear scale, where the shift transformation is not applicable or meaningful, we replace the linear transformation in Eq. 5 with a multiplication by $\tau$. For JRF, the noise term $\varepsilon_\lambda$ was always added to the log-odds of $\pi(p)$ to maintain a fair comparison between models. D1 does not influence the number of free parameters of the model.

**D2: bounded or bounds-free.** We assumed a bounding operation in BLO (Eq. 4), where probabilities outside specific boundaries are truncated to the boundaries. We considered alternative models that are bounds-free. Bounds-free models would not include the bounds parameters, $\Delta^-$ and $\Delta^+$.

**D3: variance compensation.** In BLO, $\omega_p$ is inversely related to $V(\hat{p})$ (Eq. 9) so that the encoding variance is appropriately compensated in the framework of Bayesian inference. Alternatively, we considered $\omega_p$ as a constant that does not change with $p$, as if the compensated variance is constant. When $\omega_p$ is constant, the potentially non-linear transformation of BLO (Eq. 6) is reduced to the linear transformation of LLO, as a re-parameterization would reveal (with one free parameter reduced). The two forms of variance compensation will be referred to as *V(p)* and *V=const*.

The three dimensions—D1, D2, and D3—correspond to the three assumptions of BLO. We did not list the presence or absence of the scaling factor $\tau$ ($= \frac{\Psi}{(\Delta^+ - \Delta^-)/2}$ in BLO) as a possible dimension for both theoretical and practical reasons. On one hand, $\tau$ is required to map the bounded interval to the fixed Thurstone scale. On the other hand, the absence of scaling would preclude a greater-than-one $\hat{\gamma}$. There is evidence from several laboratories other than our own experiments that $\hat{\gamma}$ can be greater than 1 (Brooke & MacRae, 1977; Pitz, 1966; Shuford, 1961; Wu, Delgado, & Maloney, 2009).

The three dimensions are independent of each other, analogous to the different factors manipulated in an experiment with a factorial design. In total, we tested 3 (D1: log-odds, Prelec, linear) × 2 (D2: bounded, bounds-free) × 2 (D3: $V(p)$, $V = const$) =

12 different models. LLO is among the 12 models (when D1=log-odds, D2=bounds-free, D3=$V = const$ ).

### Supplement S7: Additional alternative models for DMR

Among the large number of participants (75 participants) tested in Experiment JD, we noticed that a substantial minority exhibited an *S*-shaped distortion, rather than the more typical inverted-*S*-shaped distortion (Figure 2bc). We grouped the two types of participants based on their slope of distortion estimated by LLO and found that, for both DMR and JRF, the BLO model outperformed all alternative models for both types of participants (Figure S4).

All the 12 models constructed above for DMR are in the framework of Cumulative Prospect Theory (CPT). We also considered the original Prospect Theory (OPT, Kahneman & Tversky, 1979), where the probability weight for the lower outcome in the gamble, $\pi(1-p)$, is computed in the same way as $\pi(p)$, while $\pi(1-p)=1-\pi(p)$ is assumed in CPT. A parallel set of 12 models were constructed. The results of model comparison among the OPT models (Figure S5a) were similar to those of the CPT models (Figure 3a), with each OPT model fitting considerably worse than its CPT counterpart.

In addition, we considered a CPT model with Tversky and Kahneman's (1992) weighting function and two competitive models outside the framework of Prospect Theory—the transfer of attention exchange model (Birnbaum, 2005) and Haruvy's regression model (Erev et al., 2010). The transfer of attention exchange model, according to Birnbaum (2005), could outperform CPT in predicting human choices in a wide range of gambles, while Haruvy's regression model was the second best model for the decision from description dataset in Erev et al.'s (2010) model competition (the best model was a variant of CPT). However, for the DMR datasets we tested, the fits of all these additional models (see below for model definitions) were either worse than or close to the LLO model and were much worse than the BLO model (Figure S5b).

**Model definitions**

**1. CPT model with Tversky & Kahneman's (1992) weighting function.** The weighting function has the form:

$$\pi(p) = \frac{p^{\gamma}}{\left(p^{\gamma} + (1-p)^{\gamma}\right)^{\frac{1}{\gamma}}}. \tag{S14}$$

where $\gamma$ is a free parameter for the slope of probability distortion. The other assumptions of the model are identical to those of the other CPT models. In total, the model has three free parameters: $\gamma, \alpha, \sigma_{CE}^2$ .

**2. Transfer of attention exchange model (TAX, see Birnbaum, 2005).** Similar to CPT, the TAX model assumes that the weight of a branch in the computation of expected utility depends on the branch's probability and that the weights of all branches add up to 1. But different from CPT, the weight for a specific branch is also influenced by the existence of the other branches in the gamble: branches compete for attention, which results in a transfer of weight from branches with higher-valued outcomes to those with lower-valued outcomes. For an *n*-branch gamble $(x_1, p_1; x_2, p_2; ...; x_n, p_n)$ , where $x_1 > x_2 > \cdots > x_n > 0$ , the probability weight for the *i*-th branch can be written as:

$$\pi_i = \frac{t(p_i) + (\delta/(n+1))\sum_{j=1}^{i-1} t(p_j) - (\delta/(n+1))\sum_{j=i+1}^{n} t(p_j)}{\sum_{i=1}^{n} t(p_i)}, \tag{S15}$$

where $t(p) = p^{\gamma}$ , $\delta$ and $\gamma$ are free parameters. In the case of two-outcome gambles $(x_1, p; x_2, 1-p)$, Eq. S15 is reduced to:

$$\pi_1 = \frac{\left(1-\frac{\delta}{3}\right)p^\gamma}{\frac{\delta}{3}p^\gamma + (1-p)^\gamma} \quad . \qquad \text{(S16)}$$
$$\pi_2 = 1 - \pi_1$$

That is, for two-outcome gambles, TAX is mathematically equivalent to a CPT model whose weighting function is defined by Eq. S16. The assumptions of the model for utility function and noise variance are identical to those of the CPT models. In total, the model has four free parameters: $\gamma, \delta, \alpha, \sigma_{CE}^2$ .

**3. Haruvy's regression model (Erev et al., 2010).** Haruvy's regression model treats the prediction of human choices as a regression problem. It was originally a logistic regression model designed for the forced-choice task between a two-outcome gamble and a sure payoff. For gamble $G = (x_1, p; x_2, 1-p)$ and sure payoff $y$ , it defines the tendency to prefer the gamble as

$$T(G) = \beta_0 + \beta_1 x_1 + \beta_2 x_2 + \beta_3 y + \gamma_1 p + \gamma_2 EV + \gamma_3 I \ , \qquad \text{(S17)}$$

where $EV = px_1 + (1-p)x_2$ is the expected value of the gamble, $I$ is an indicator variable that equals 1 if $EV > y$ and 0 otherwise, $\beta_0, \beta_1, \beta_2, \beta_3, \gamma_1, \gamma_2, \gamma_3$ are free parameters. The probability of choosing the gamble is modeled as a logistic function of $T(G)$:

$$\Pr(G) = \frac{1}{1+e^{-T(G)}} \ . \qquad \text{(S18)}$$

For the DMR task in the current paper, certainty equivalent (CE) is measured for each gamble. To apply Haruvy's regression model, we consider the problem as choosing between a gamble and its CE. The definition of CE implies that $\Pr(G) = 0.5$, which requires $T(G) = 0$. Thus the goal is to fit the following linear regression model:

$$0 = \beta_0 + \beta_1 x_1 + \beta_2 x_2 + \beta_3 CE + \gamma_1 p + \gamma_2 EV + \gamma_3 I + Normal\left(0, \sigma_{CE}^2\right). \quad \text{(S19)}$$

It is obvious that one of the parameters in Eq. S19 is redundant. We arbitrarily set $\beta_3 = -1$. In total, the model has seven free parameters: $\beta_0, \beta_1, \beta_2, \gamma_1, \gamma_2, \gamma_3, \sigma_{CE}^2$ .

### Supplement S8: Additional alternative models and datasets for JRF

For the JRF task, we assumed additional noise from sampling without replacement. An alternative assumption is sampling with replacement, where the variance of sampling error is

$$V(\hat{p}) = \frac{p(1-p)}{n_s} \tag{S20}$$

For $\omega_p = \frac{1}{1+\kappa V(\hat{p})}$ , the $n_s$ would be assimilated into $\kappa$ , and we can simply write

$$V(\hat{p}) \propto p(1-p) \ , \tag{S21}$$

which is identical to what we have for DMR based on the assumption of Gaussian noise on the Thurston scale. In the framework of factorial model comparison, for JRF we constructed 6 additional models all of which assume sampling with replacement for variance compensation but differ in the scale of transformation (log-odds, Prelec, or linear) and boundedness (bounded or bounds-free). We fitted these models to the 75 participants of Experiment JD and presented the extended results of factorial model comparison (12 + 6 = 18 models) in Figure S9a. Models assuming sampling with replacement performed worse than those of sampling without replacement.

We also re-analyzed a JRF dataset from Zhang and Maloney's (2012) Experiment 1, which used similar task and design as the JRF task of Experiment JD except that the value of relative frequency ranged from 0.01 to 0.99 in steps of 0.01. Its results of factorial model comparison (Figure S9b) were similar to those of Experiment JD. The BLO model assuming sampling without replacement was still the best fitting model.

Zhang and Maloney (2012) also reported a numerosity effect (see their Figure 8): the slope of distortion, $\hat{\gamma}$, decreases with the numerosity (total number of dots) of the display. As shown in Figure S10, the BLO model (assuming sampling without replacement) could well predict the numerosity effect of slope. BLO could even predict the slight increase of the crossover point, $\hat{p}_0$, with the numerosity.

### Supplement S9: Analysis of Individual Parameter Settings

The estimated parameters of BLO, summarized by their median values across participants, are summarized in Supplemental Table S2. As described in the main text (Figure 1b), the lower bound $\Delta^-$ and upper bound $\Delta^+$ define the range of input in log-odds that the brain chooses to represent at the moment. Any value outside the range cannot be precisely represented but cut off to its closest boundary. The parameter $\Psi$ is the half-range of the Thurstone scale. The anchor $\Lambda_0$ is the fixed point on the log-odds scale to expand or contract around in the compensation of encoding variance. The parameter $\kappa$ controls the extent to which the encoding uncertainty influences the weight $\omega_p$ for bounded log-odds.

What parameters of probability distortion may be relatively invariant across tasks and time? For Experiment JD where participants completed two different tasks or even two sessions, we computed Spearman's correlations across tasks and across sessions for each BLO or LLO parameter that is shared by DMR and JRF, and for measures derived from BLO parameters, including $\Delta \equiv \left(\Delta^+ - \Delta^-\right)/2$ , $\tau \equiv \Psi/\Delta$ , and $\Psi/\sigma_\lambda$ or $\Psi/\sigma_{CE}$ (as proxy for the Thurstone capacity $\Psi/\sigma_\Psi$ ). The results are shown in Supplemental Table S3, with $r_s$ referring to Spearman's correlation coefficient and the $P$ value being right-tailed. Among these 12 measures of probability distortion, $\Psi$ and $\Psi/\sigma_{\lambda/CE}$ were the only two measures whose across-task and across-session correlations were all significant.

### Supplement S10: Computation of mutual information or expected error

For a specific real or virtual participant in a specific task, we used the BLO model to randomly generate simulated subjective estimates for the objective probabilities of the experiment to compute the expected mutual information or the expected error (root of mean squared error) between objective and subjective probabilities, as defined in the main text. To obtain a stable estimate of the expected mutual information or expected error, we repeated the stimulus set of each task to produce 198,000 trials. In the numerical computation of mutual information, the objective and subjective probabilities were quantized by rounding to the 2nd decimal.

For a specific real or virtual participant, the standard deviation of the Gaussian noise *on the Thurstone scale*, $\sigma_\Psi$, is constant for different bounds parameters (as illustrated in Figure 1b). See Supplement S11 for how $\sigma_\Psi$ was estimated from participants' BLO parameters.

When Weber fraction $f_w > 0$, we first generated $\lambda\left[\hat{\pi}(p)\right]$ that is perturbed by Gaussian noise on the Thurstone scale, transformed into $\hat{\pi}(p)$, and then applied additional multiplicative Gaussian random noise (standard deviation $f_w\hat{\pi}(p)$) to $\hat{\pi}(p)$. The final estimate of probability was forced to be between 0 and 1.

For DMR, the deviation of the observed $\left[\Delta^-,\Delta^+\right]$ from optimality in expected mutual information is a *U*-shaped function of $f_w$, which would come close to 0 for some medium level of $f_w$. We grid searched (in steps of 0.01) the $f_w$ that minimizes the deviation between the observed and optimal $\left[\Delta^-,\Delta^+\right]$ in expected mutual information and found $f_w$ =0.19. The contour of expected mutual information for $f_w$ =0.19 is plot in Figure 5c.

For the bounds-free representation in Figure 5c, we assume that no bounds had been imposed on the probability range of $[0.01, 0.99]$, that is, setting $\Delta^- = -4.6$ and $\Delta^+ = 4.6$.

For the no-variance-compensation representation in Figure 6c, we assume that the noise-perturbed $\Gamma(\lambda[p])$, without any scaling or variance compensation, had been transformed into probability as the subjective estimate of probability.

### Supplement S11: Estimating noise SD on the Thurstone scale

The standard deviation (SD) of the noise on the bounded Thurstone scale $[-\Psi, \Psi]$ is denoted $\sigma_\Psi$, as in our definition of the Thurstone capacity $\Psi/\sigma_\Psi$. We estimated participants' $\sigma_\Psi$ separately for JRF and DMR using the procedure described below.

In fitting BLO to JRF data, we estimated the noise SD in participants' subjective log-odds, $\sigma_\lambda$ (as in Eq. 10), which is different from $\sigma_\Psi$ by a factor of $\omega_p$ according to the variance compensation specified in Eq. 6. Because $\omega_p$ is not constant but varies with *p*, we could not just divide $\sigma_\lambda$ by $\omega_p$ to obtain an estimate of $\sigma_\Psi$. Instead, we used the slope of distortion, $\gamma$, of LLO as an estimate of the average $\tau\omega_p$ (following Eq. S7), and thus had $\sigma_\Psi^{JRF} \approx \frac{\tau^{JRF}}{\gamma^{JRF}} \sigma_\lambda^{JRF}$.

For DMR, where the noise in the subjective log-odds could not be directly estimated, we estimated the noise SD on the Thurstone scale based on the assumption of equivalence of Thurstone capacity across tasks: $\Psi^{JRF}/\sigma_\Psi^{JRF} = \Psi^{DMR}/\sigma_\Psi^{DMR}$. That is, $\sigma_\Psi^{DMR} = \frac{\Psi^{DMR}}{\Psi^{JRF}} \sigma_\Psi^{JRF}$. We then estimated the noise SD in the subjective log-odds to be $\sigma_\lambda^{DMR} \approx \frac{\gamma^{DMR}}{\tau^{DMR}} \sigma_\Psi^{DMR}$.

These estimates in noise SD were used in generating the simulated responses needed for computing mutual information or expected error. We must caution that some simplified assumptions have been taken to make these computations feasible. For example, in the JRF task, the noises in the encoded log-odds include the Gaussian noise on the Thurstone scale as well as the random sampling error, but we did not treat them separately.

### Supplement S12: BLO and optimal variance compensation

As described in the main text, for any probability *p*, BLO assumes the following transformation on the log-odds scale:

$$\Lambda^{\omega}(p)=\omega_p\tau\left[\Gamma\left(\lambda\left[p\right]\right)-\left(\Delta^{-}+\Delta^{+}\right)/2\right]+\left(1-\omega_p\right)\Lambda_0\,, \tag{S22}$$

where the scaling factor $\tau$ is equivalent to $\frac{\Psi}{\left(\Delta^{+}-\Delta^{-}\right)/2}$ in Eq. 5 (i.e. $\tau\equiv\Psi/\Delta$ ). The reliability parameter $\omega_p$ that compensates for encoding variance varies with *p* and has the particular form

$$\omega_p=\frac{1}{1+\kappa p\left(1-p\right)} \tag{S23}$$

for DMR and a similar (proportional to *p(1–p)*) form for JRF. Compared with some classic examples of variance compensation such as cue combination (Oruç, Maloney, & Landy, 2003), the transformation specified by Eq. S22 seems to be unusual in two aspects. First, the reliability parameter $\omega_p$ is not constant but varies with *p*. Second, there is a seemingly unnecessary multiplication operation by $\tau$ , whose value was estimated to be greater than one in both DMR and JRF. However, we will demonstrate that both the varying $\omega_p$ and greater-than-one $\tau$ in Eq. S22 are necessary, if our goal is to minimize the expected deviation between objective and subjective probabilities. Eq. S22 differs from the classic variance compensation solution based on Gaussian prior and likelihood because of the log-odds and truncation transformations involved in our problem.

For simplicity, we first omit the truncation transformation, $\Gamma(\bullet)$ , and frame variance compensation as the following Bayesian decision problem: Given the noisy estimate $\hat{p}$ (or equivalently, the value of $\lambda(\hat{p})$), what should one conclude about the likely true probability $p$? More precisely, we ask, given the likelihood function $L\left[\lambda(p)\middle|\lambda(\hat{p})\right]$ and the prior distribution $\Pi\left[\lambda(p)\right]$, what posterior estimate of $p$ (denoted $p^{post}$) would minimize the expected squared error between $p^{post}$ and $p$?

According to Bayes' theorem, given the noisy observation $\lambda(\hat{p})$, the posterior distribution of $\lambda(p)$ is

$$\Pr\left[\lambda(p)\middle|\lambda(\hat{p})\right] \propto \Pi\left[\lambda(p)\right] L\left[\lambda(p)\middle|\lambda(\hat{p})\right]. \tag{S24}$$

Note that the posterior distribution we finally need for the Bayesian decision problem above is not $\Pr\left[\lambda(p)\middle|\lambda(\hat{p})\right]$, but $\Pr\left[p\middle|\lambda(\hat{p})\right]$. Following the transformation rule of probability distributions (Wasserman, 2004), $f(x) = g(y)\dfrac{dy}{dx}$ , we have:

$$\begin{aligned} \Pr\left[p\middle|\lambda(\hat{p})\right] &= \Pr\left[\lambda(p)\middle|\lambda(\hat{p})\right]\lambda'(p) \\ &= \frac{1}{p(1-p)}\Pr\left[\lambda(p)\middle|\lambda(\hat{p})\right] \end{aligned}. \tag{S25}$$

Denote an estimate of $p$ based on the observed $\lambda(\hat{p})$ as $\eta(\hat{p})$. The optimal estimate $p^{post}$ is the function $\eta(\hat{p})$ that minimizes the expected squared error in probability:

$$\begin{aligned} p^{post} &= \operatorname{argmin}_{\eta(\hat{p})} E_{p|\lambda(\hat{p})}\left[\left(\eta(\hat{p}) - p\right)^2\right] \\ &= \operatorname{argmin}_{\eta(\hat{p})} \int_p \left(\eta(\hat{p}) - p\right)^2 \Pr\left[p\middle|\lambda(\hat{p})\right] dp \end{aligned}. \tag{S26}$$

If the optimization problem had been defined for the log-odds scale, the Gaussian prior and likelihood on the log-odds scale would have implied a Gaussian posterior distribution, which corresponds to a simple analytical solution for variance compensation (Oruç, Maloney, & Landy, 2003). However, the posterior distribution for our optimization problem on the probability scale, $\Pr\left[p\middle|\lambda(\hat{p})\right]$, is *not* a Gaussian distribution and its optimal estimate $p^{post}$ has no known analytical form.

The problem gets even complicated when we take into account the truncation operation $\Gamma(\bullet)$ imposed on $\lambda(\hat{p})$. The truncation operation, which is a function of the bounds parameters $\Delta^-$ and $\Delta^+$, makes it impossible to fully recover the objective probabilities outside the bounds and also changes the optimal decoding rule. That is, reverting the linear mapping from $\left[\Delta^-,\Delta^+\right]$ to $\left[-\Psi,\Psi\right]$, or in terms of our previous parameterization, dividing out the scaling factor $\tau$ ($\equiv \Psi/\Delta$), does not necessarily be part of the optimal decoding rule that minimizes expected error in subjective probability.

Though there is no analytical form for the optimal decoding rule, we performed a numerical simulation to compare the effect of $\tau = \Psi/\Delta$ with $\tau = 1$, and the effect of $\omega_p = \dfrac{1}{1+\kappa p(1-p)}$ with constant $\omega$ in reducing the expected error of the subjective estimate of probability. In our simulations, we considered four different prior distributions of objective log-odds and three different noise levels based on the stimuli used in previous decision under risk studies and the parameters of probability distortion estimated in Experiment JD. To facilitate the comparison between $\omega_p$ and $\omega$, we write $\omega = \dfrac{1}{1+\kappa C}$, where $C$ denotes the median of *p(1–p)* for a specific distribution of objective probabilities. For each prior distribution and noise level, we generated 99,000 samples of $p$ and $\lambda(\hat{p})$, and computed the expected error as a function of $\kappa$ for different combinations of $\tau$ and $\omega_p$. As Figure S8 shows, the minimum expected error for $\tau = \Psi/\Delta$ is smaller than the minimum expected error for $\tau = 1$, and the minimum expected error for $\omega_p$ is smaller than the minimum expected error for constant $\omega$.

### Supplement S13: Predicted violations of stochastic dominance

We would like to highlight a counterintuitive prediction of BLO that agrees with empirical observations, the *violation of stochastic dominance* observed for one participant in the DMR experiment of GW99. A probability weighting function, $\pi(p)$, predicted by LLO would be monotonically increasing with the objective probability, but that of BLO need not be, as demonstrated by the $\pi(p)$ of Participant 4 in GW99 (see the fourth panel in Figure 2a). The possible non-monotonicity comes from the non-linear transformation that compensates for encoding variance, where the weight for the anchor depends on the value of the objective probability, and may be exaggerated by boundedness. At first sign this might sound like a "bug" in the assumptions of BLO—the interaction of bounding and anchoring operations can, in theory, result in a non-monotonic $\pi(p)$ (the $\pi(p)$ predicted by the bounded Prelec model had similar non-monotonicity). To our surprise, a closer examination of Participant 4's estimated CE did provide evidence for violation of stochastic dominance: For two lotteries $(x_1, p; x_2, 1-p)$, where $x_1 > x_2 \geq 0$ are identical, stochastic dominance predicts that the CE for a smaller value of *p* should be smaller than the CE for a larger value of $p$. Among the probabilities used in the experiment, the value *p* = 0.25 is close to the local maximum and the values *p* = 0.40, 0.50, or 0.60 are close to the local minimum. As a consequence of the non-monotonicity of the probability distortion function for the anomalous subject, $\pi(0.25)$ is higher than $\pi(0.40)$, $\pi(0.50)$, or $\pi(0.60)$, we may expect violations of stochastic dominance, if any, to occur in the comparisons of the CE of *p* = 0.25 to the CE of *p* = 0.40, 0.50, or 0.60. For Participant 4, among the 45 pairs of CE comparisons, the CE of *p* = 0.25 was *larger* 29 times (a match counts as 0.5 times). According to Fisher's exact test, we could reject the null hypothesis that the CE of *p* = 0.25 is larger in no more than half of the pairs at a significance level of

0.036. That is, for the participant, the observed non-monotonicity in $\pi(p)$ agrees with her consistent violation of stochastic dominance in specific risky choices.

The violation of stochastic dominance in Participant 4's choice behaviors was subtle, going unnoticed in the original paper (Gonzalez & Wu, 1999). We noticed it only because the BLO model fitted to this participant predicted such violations. There may well be more such participants overlooked in previous studies.

The BLO model also permits a more dramatic form of violation of stochastic dominance, one that is also reported in the literature: given a choice between a sure reward and a lottery with probability $p < 1$ of receiving the same reward and otherwise nothing, some participants choose the lottery more than 50% of the time (Blankenstein, Crone, van den Bos, & van Duijvenvoorde, 2016; Tymula et al., 2012). Their choices implied that $\pi(1) < \pi(p)$, that is, $\pi(p)$ is not an increasing (more precisely, non-decreasing) function of $p$. In past work, such participants were typically excluded from further analyses. As we saw above, these participants' violation of stochastic dominance is predicted by BLO for certain parameter settings that result in a non-monotonic $\pi(p)$.

Birnbaum and colleagues' elegant transfer-of-attention-exchange model (Birnbaum, 2005; Birnbaum & Navarrete, 1998) predicted the violations of stochastic dominance on the group level in a series of gambles. However, for gambles with only two branches, the model is reduced to a variant of CPT whose probability weighting function $\pi(p)$ varies monotonically with $p$ (Supplement S7, Eq. S16). As the result, it cannot explain the violation of stochastic dominance in our case. In other words, the BLO model and the transfer-of-attention-exchange model account for violations of stochastic dominance under different circumstances.

### Supplement S14: Using BLO to predict probability distortions in decision under risk

We found that subject to the dynamic range limit of representing probability, participants' choice of bounds parameters was close to maximizing the mutual information and their choice of variance compensation parameters was close to minimizing the expected deviation between objective and subjective probabilities. It implies that participants' distortion of probability may change with the distribution of objective probabilities.

Based on the observed constraint (Thurstone capacity) and bounded rationality, we predicted the probability distortion for each of the 12 DMR studies in our meta-analysis (see Table S4), where different distributions of objective probabilities were used. In particular, we assumed that the $\Psi$ and $\sigma_{\Psi}$ of each study were the same as the median values estimated for the 75 participants in our DMR task (see Supplement S11 for the estimation of $\sigma_{\Psi}$). We assumed that the $\Lambda_0$ of each study equals the mean of the objective log-odds in the study. For each study, we first set $\kappa$ to be the median estimated from our 75 participants and used numerical simulations to find the $\Delta^-$ and $\Delta^+$ that maximize expected mutual information. Based on the optimal $\Delta^-$ and $\Delta^+$, we then searched for the optimal $\kappa$ that minimizes expected error. The Weber fraction of the additional multiplicative noise was set to be 0.19, as that best explained the observed $\left(\Delta^-,\Delta^+\right)$ (see Figure 5c). Finally, we used BLO models with the optimal $\Delta^-$ and $\Delta^+$, and the optimal $\kappa$ to generate simulated subjective probabilities and estimated the slope of distortion from LLO fits.

### Supplemental Tables

Table S1. *Notations*

| | |
|---|---|
| **General** | |
| $p$ | A generic value on the probability scale |
| $\lambda$ | A generic value on the log-odds scale |
| $\lambda(p)$ | Log-odds of probability *p*. $\lambda(p)=\log(p/(1-p))$ |
| $\pi(p)$ | Distorted probability (also denoted as $w(p)$, the decision weight, in decision tasks) |
| **Linear in Log-Odds (LLO) Model** | |
| $\gamma$ | Slope of the observed linear transformation of log odds |
| $p_0$ | Crossover point; controlling the intercept of the linear transformation of log odds |
| **Bounded Log-Odds (BLO) Model** | |
| $\Delta^-,\Delta^+$ | Controlling the bounds of the log-odds representation |
| $\Psi$ | Half-range of the Thurstone scale |
| $\Lambda_0$ | Anchor parameter |
| $\kappa$ | Parameter that controls the extent to which encoding variance influences $\omega_p$ |
| $V(\hat{p})$ | Encoding variance of the estimate $\hat{p}$ |
| $\omega_p$ | Reliability parameter that is inversely related to encoding variance |
| $\Gamma(\bullet)$ | Bounding operation |
| $\Delta\equiv(\Delta^+-\Delta^-)/2$ | Half-range of the encoded bounded interval |
| $\tau\equiv\Psi/\Delta$ | Scaling parameter |
| $\Psi/\sigma_\Psi$ | Thurstone capacity |
| **Alternative Models** | |
| $\lambda^I(p)$ | Converting *p* to the Prelec scale |
| $\lambda^O(p)$ | Converting *p* to the linear scale |
| **Judgment of Relative Frequency (JRF)** | |
| $n_s$ | Sample size; controlling $V(\hat{p})$ |
| $\sigma_\lambda^2$ | Noise variance of judgment on the log-odds scale |
| $p$ | Relative frequency of black or white dots in the array |
| **Decision-Making under Risk (DMR)** | |
| $\alpha$ | Exponent of the power utility function |
| $\sigma_{CE}^2$ | Noise variance of certainty equivalent |
| $p$ | Probability of a specific outcome in a lottery |

Table S2. *Estimated parameters of BLO*

| | Bounds | | Thurstone half-range | Variance compensation | | Noise | Sample size (JRF) | Value (DMR) |
|---|---|---|---|---|---|---|---|---|
| **Experiment** | $\Delta^-$ | $\Delta^+$ | $\Psi$ | $\Lambda_0$ | $\kappa$ | $\sigma_\lambda^2$ or $\sigma_{CE}^2$ | $n_s$ | $\alpha$ |
| GW99 (DMR) | –0.12 (0.47) | 1.74 (0.85) | 1.78 | 0.13 (0.53) | 5.91 | 385.6 [ξ] | N/A | 0.52 |
| JD, DMR | –0.60 (0.35) | 1.75 (0.85) | 2.04 | –0.05 (0.49) | 5.20 | 474.6 [ξ] | N/A | 0.72 |
| JD, JRF | –1.64 (0.16) | 1.38 (0.80) | 3.90 | 0.03 (0.51) | 343.4 | 0.45 | 37.1 | N/A |
| ZM12 (JRF) | –1.13 (0.25) | 1.34 (0.79) | 3.58 | 0.15 (0.54) | 1023 | 0.42 | 93.9 | N/A |

*Note*. Medians across participants are reported. Value inside the parentheses is the probability equivalent of the parameter (originally on the log-odds scale).

[ξ] The noise parameter for DMR is denoted $\sigma_{CE}^2$, which is defined on a different scale than the $\sigma_\lambda^2$ of JRF.

Table S3. *Across-task and across-session correlations for BLO- and LLO-derived measures*

| Measures | Across-task correlation | | Across-session correlation in DMR | | Across-session correlation in JRF | |
|---|---|---|---|---|---|---|
| | $r_s$ | $P$ | $r_s$ | $P$ | $r_s$ | $P$ |
| BLO | | | | | | |
| $\Delta^-$ | 0.068 | 0.28 | *0.36* | *0.004* | *0.43* | *< 0.001* |
| $\Delta^+$ | –0.012 | 0.54 | 0.13 | 0.18 | *0.31* | *0.013* |
| $\Psi$ | ***0.23*** | ***0.026*** | ***0.57*** | ***< 0.001*** | ***0.83*** | ***< 0.001*** |
| $\Lambda_0$ | 0.075 | 0.26 | –0.006 | 0.52 | *0.39* | *0.003* |
| $\kappa$ | –0.11 | 0.82 | 0.18 | 0.10 | *0.72* | *< 0.001* |
| $\sigma_\lambda$ or $\sigma_{CE}$ | 0.069 | 0.28 | *0.49* | *< 0.001* | *0.55* | *< 0.001* |
| $\Delta \equiv (\Delta^+ - \Delta^-)/2$ | –0.035 | 0.62 | 0.017 | 0.45 | *0.36* | *< 0.001* |
| $\tau \equiv \Psi/\Delta$ | –0.027 | 0.59 | –0.28 | 0.98 | *0.39* | *0.003* |
| $\Psi/\sigma_{\lambda/CE}$ | ***0.40*** | ***< 0.001*** | ***0.60*** | ***< 0.001*** | ***0.56*** | ***< 0.001*** |
| LLO | | | | | | |
| $\gamma$ | 0.10 | 0.20 | 0.67 | < 0.001 | *0.85* | *< 0.001* |
| $p_0$ | *0.23* | *0.025* | 0.041 | 0.39 | *0.60* | *< 0.001* |
| $\sigma_\lambda$ or $\sigma_{CE}$ | 0.011 | 0.46 | 0.42 | < 0.001 | *0.57* | *< 0.001* |

*Note*. Across-task correlations were computed for the 75 participants of Experiment JD, who completed both the DMR and JRF tasks. Across-session correlations were computed for the 51 participants who had completed two sessions on two different days. The $r_s$ refers to Spearman's correlation coefficient and the *P* value is right-tailed. Significant positive correlations ($P$ < 0.05) are highlighted in Italic. Measures whose across-task and across-session correlations were all significant are highlighted in bold.

Table S4. *Studies used in the meta-analysis on the slope of probability distortion in decision under risk*

| **Study** | $\gamma$ | **Distribution of *p* in** $(x_1, p; x_2, 1-p)$ |
|---|---|---|
| Abdellaoui, Diecidue, and Öncüler (2011) | 0.62 (W3A) | 1/6, 2/6, 2/6, 3/6, 4/6, 5/6 |
| Abdellaoui, L'Haridon, and Paraschiv (2011) | 0.65 (W2) | 0.05, 0.25, 0.5, 0.75, 0.95 |
| Bruhin, Fehr-Duda, and Epper (2010) | $0.42^{\xi}$ (W2) | 0.05×4, 0.1×3, 0.25×3, 0.5×6, 0.75×3, 0.9×3, 0.95×3 |
| Glöckner and Pachur (2012) | 0.61 (W1)<br>0.67 (W2) | 180 gambles from Rieskamp (2008);<br>40 gambles from Glöckner and Betsch (2008);<br>36 gambles from Holt and Laury (2002) and Gächter, Johnson, and Herrmann (2007) |
| Gonzalez and Wu (1999) | 0.44 (W2) | 0.01, 0.05, 0.1, 0.25, 0.4, 0.5, 0.6, 0.75, 0.9, 0.95, 0.99 |
| Harrison and Rutström (2009) | 0.91 (W1) | 0.13, 0.25, 0.37, 0.5, 0.62, 0.75, 0.87 |
| Rieskamp (2008) | 0.77 (W1) | 0.01, 0.02, …, 0.99 |
| Stott (2006) | 0.96 (W1)<br>0.96 (W2)<br>1.00 (W3A)<br>0.94 (W3B) | 0.1×34, 0.2×14, 0.3×20, 0.4×20, 0.5×21, 0.6×18, 0.7×12, 0.8×14, 0.9×27 |
| Tanaka, Camerer, and Nguyen (2010) | 0.74 (W3B) | 0.1, 0.3, 0.7, 0.9 |
| Tversky and Kahneman (1992) | 0.61 (W1) | 0.01×2, 0.05×3, 0.1×3, 0.25×3, 0.5×6, 0.75×3, 0.9×3, 0.95×3, 0.99×2 |
| Vrecko and Langer (2013) | 0.72 (W1) | Same as Zeisberger et al., 2012 (see below) |
| Zeisberger, Vrecko, and Langer (2012) | 0.86 (W1) | 0.2×6, 0.5×12, 0.8×6 |

*Note*. All studies were selected from Table A.3 of Fox and Poldrack (2014) using the criteria described in the text. The $\gamma$ column corresponds to $\gamma^+$ in Fox and Poldrack (2014), with the codes in the parentheses denoting the functional form of the probability weighting function. Following Fox and Poldrack (2014), "W1" refers to Tversky and

Kahneman's (1992) one-parameter form, "W2" refers to LLO, "W3A" and "W3B" refer respectively to two- and one-parameter Prelec functions. In the column of distribution of *p*, "*q*×*f*" (e.g. 0.05×4) denotes that the probability *q* is presented for *f* times (relative, not absolute count) in the gambles of the study.

Among these studies a few provided slope parameters estimated from multiple types of weighting functions: 0.67 and 0.61 for Glöckner and Pachur (2012), 0.96, 1.00, 0.94, and 0.96 for Stott (2006). We can see the slope parameters estimated from different types of probability weighting functions were almost the same. In our analysis, when multiple slope parameters had been reported for a study, we used their mean. Since not all studies involved the loss domain, we only considered the slope of distortion for the gain domain, that is, the $\gamma^{+}$ in Fox and Poldrack (2014).

$^{\xi}$ The value of $\gamma$ listed here (0.42) is the result of Bruhin et al.'s (2010) ZH03 experiment, which is slightly different from the pooled value in Fox and Poldrack' (2014) Table A.3 (0.38). We could only use the ZH03 experiment because Bruhin et al. (2010) did not provide the full gamble set for their other experiments that we would need for our analysis.

### Supplemental Figures

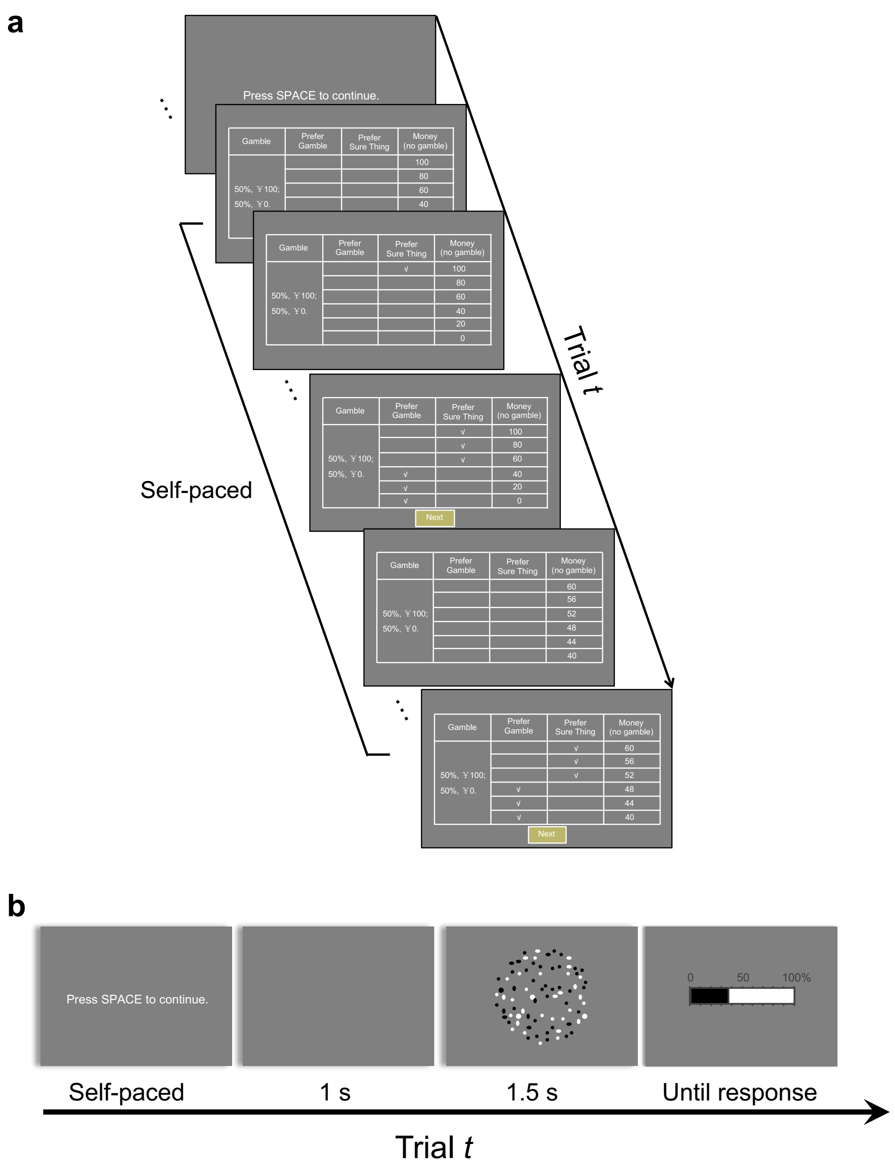


**Figure S1. The tasks. a. Decision-Making under Risk (DMR).** The procedure is based on that of Gonzalez and Wu (1999). On each trial, participants were presented with a two-outcome gamble $(x_1, p; x_2, 1-p)$ and tables of sure amounts of rewards. They were asked to check on each row of the tables whether they preferred the gamble or the sure amount. The range of the sure amounts started with $[x_2, x_1]$, and was narrowed down in the second table so that we could estimate participants' certainty equivalent (CE) for the gamble. **b. Judgment of Relative Frequency (JRF).** The procedure is based on that of Zhang and Maloney (2012). On each trial, participants were presented with an array of black and white dots and reported their estimate of the relative-frequency of black or white dots by clicking on a horizontal bar with tick marks from 0 to 100%.

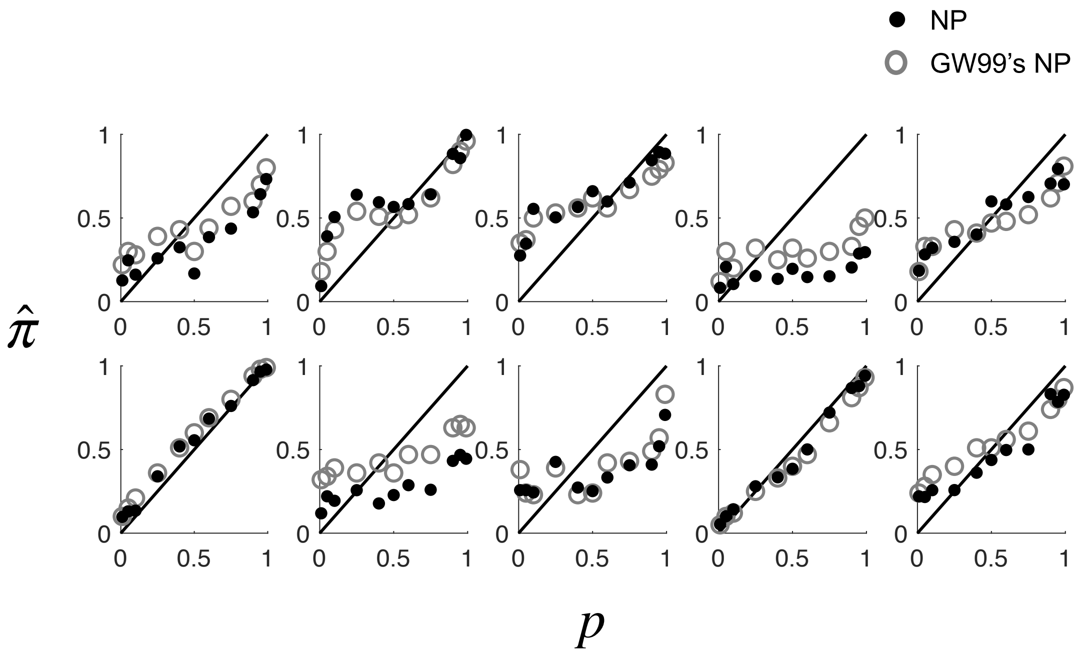


**Figure S2. Our non-parametric estimates versus Gonzalez and Wu's (1999) original non-parametric estimates of probability distortions for their 10 participants.** Our estimation procedures differed from theirs in that we assumed no functional form for the probability weighting function but a power function for the utility function, while they assumed functional forms for neither. The results from the two procedures were similar.

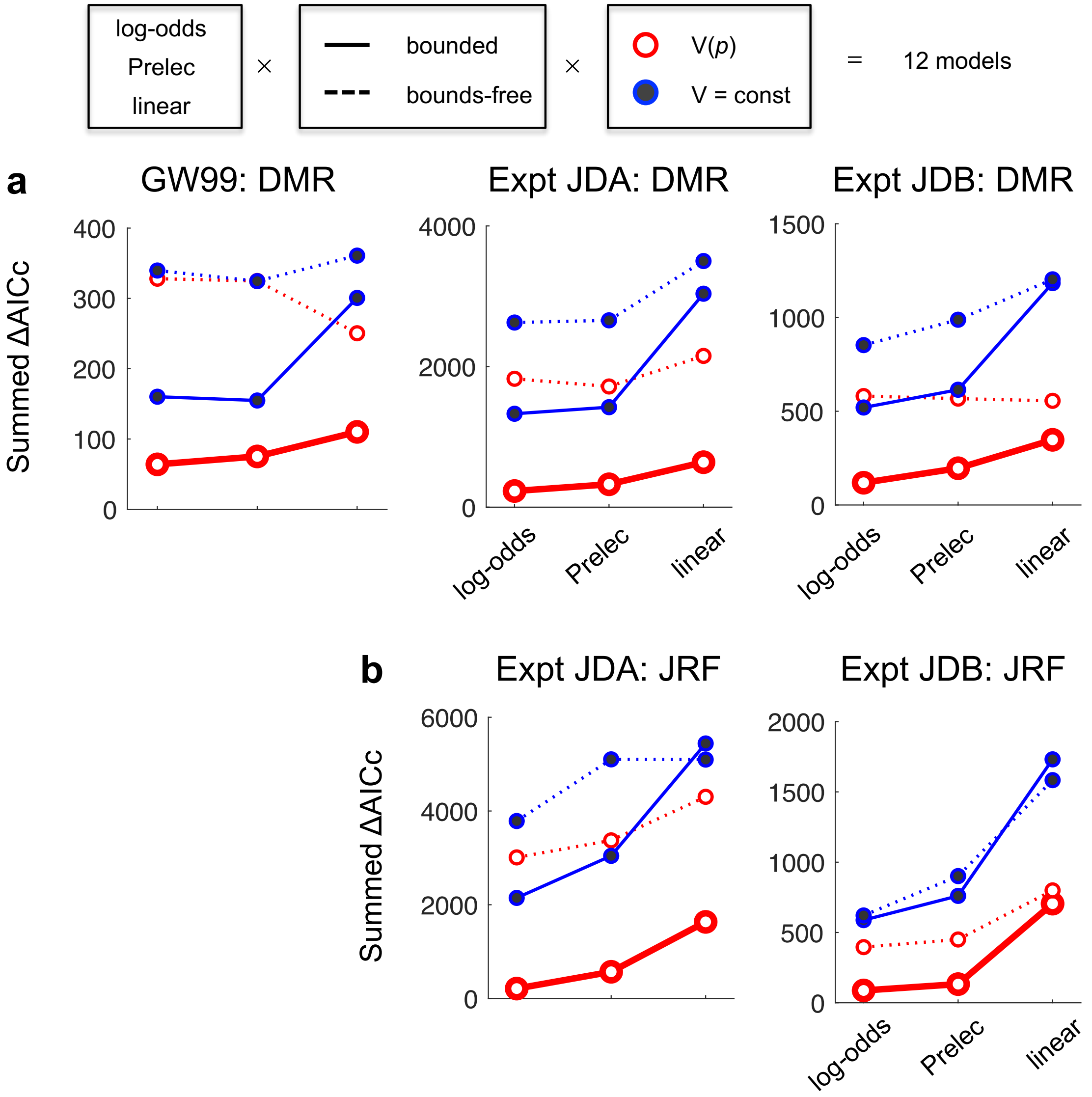


**Figure S3. Results of factorial model comparison separately for the datasets of Gonzalez and Wu (1999), Experiment JDA and JDB.** The results were similar across different datasets.

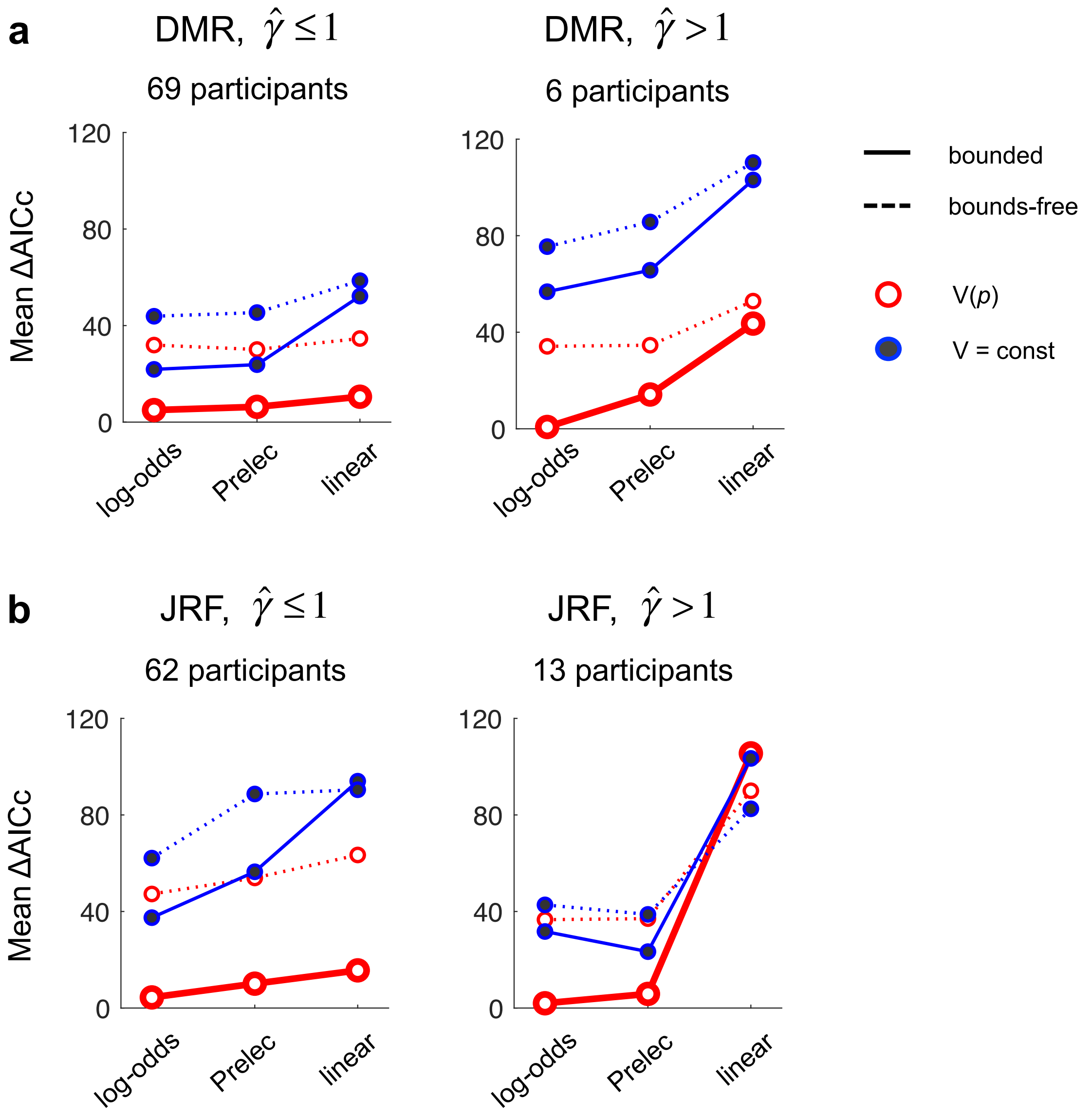


**Figure S4. Results of factorial model comparison separately for participants of $\hat{\gamma} \le 1$ and $\hat{\gamma} > 1$. a.** The JRF task. **b.** The DMR task. The mean ΔAICc across participants is plotted for each model and task. The $\hat{\gamma}$ refers to the estimated slope parameter of the LLO model. The horizontal axis corresponds to D1, scale of transformation. Solid and dotted lines code alternative models on D2, boundedness. Different colors code alternative models on D3, variance compensation. No matter whether $\hat{\gamma} \le 1$ or $\hat{\gamma} > 1$, the BLO model outperformed the alternative models.

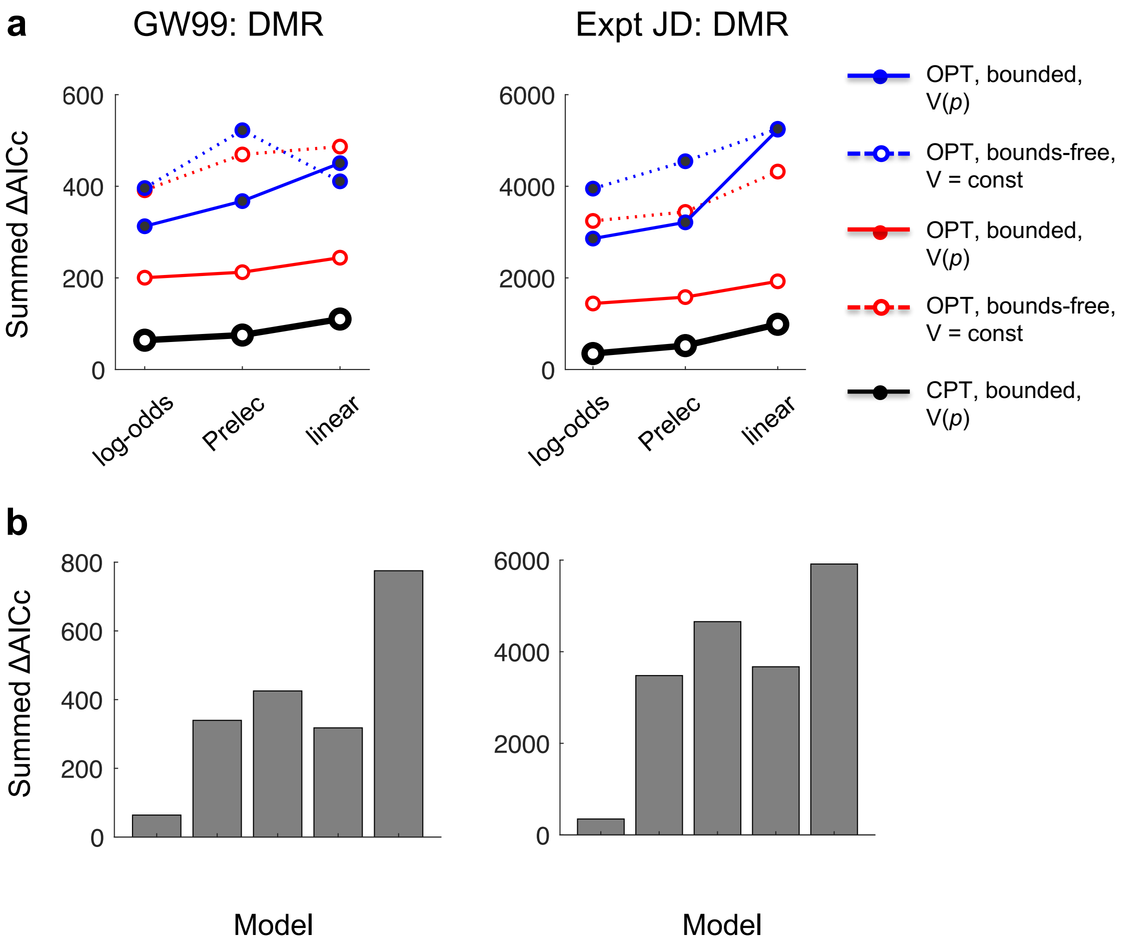


**Figure S5. Model comparison results of BLO versus additional alternative models for DMR, separately for Gonzalez and Wu' (1999) dataset (GW99) and our experiment (JD).** **a.** Models in the framework of the original Prospect Theory (OPT). The goodness-of-fit of OPT models were worse than their counterpart CPT (Cumulative Prospect Theory) models. **b.** Models of alternative form of probability weighting function or outside the framework of Prospect Theory. TK = Tversky & Kahneman's (1992) weighting function, in the CPT framework. TAX = Transfer of Attention Exchange model (Birnbaum, 2005). HR = Haruvy's regression model (Erev et al., 2010). These alternative models fit either much worse than LLO (HR), or comparable to LLO (TK, TAX), but still much worse than BLO.

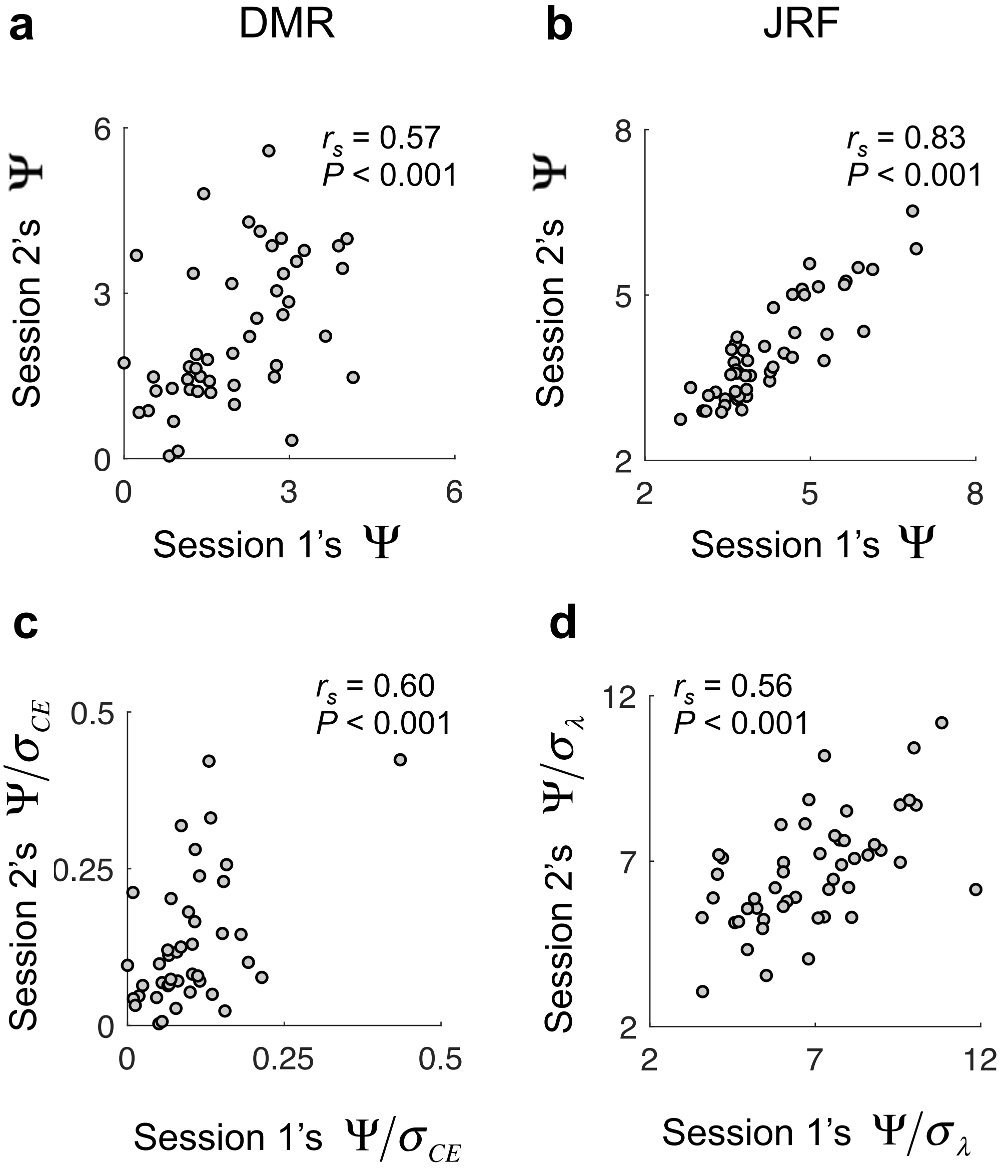


**Figure S6. Correlation of $\Psi$ and $\Psi/\sigma_{\lambda/CE}$ across time.** The correlations were based on the BLO fits of Experiment JDA (51 participants), where participants completed two sessions on two different days. Each circle is for one participant. **a.** Across-session correlation in $\Psi$ for DMR. (4/51 data points are outside the plot range.) **b.** Across-session correlation in $\Psi$ for JRF. (1/51 data points are outside the plot range.) **c.** Across-session correlation in $\Psi/\sigma_{CE}$ for DMR. (9/51 data points are outside the plot range.) **d.** Across-session correlation in $\Psi/\sigma_{\lambda}$ for JRF. (2/51 data points are outside the plot range.) The $r_s$ on the plot refers to Spearman's correlation coefficient, which is robust to outliers, and $P$ is right-tailed.

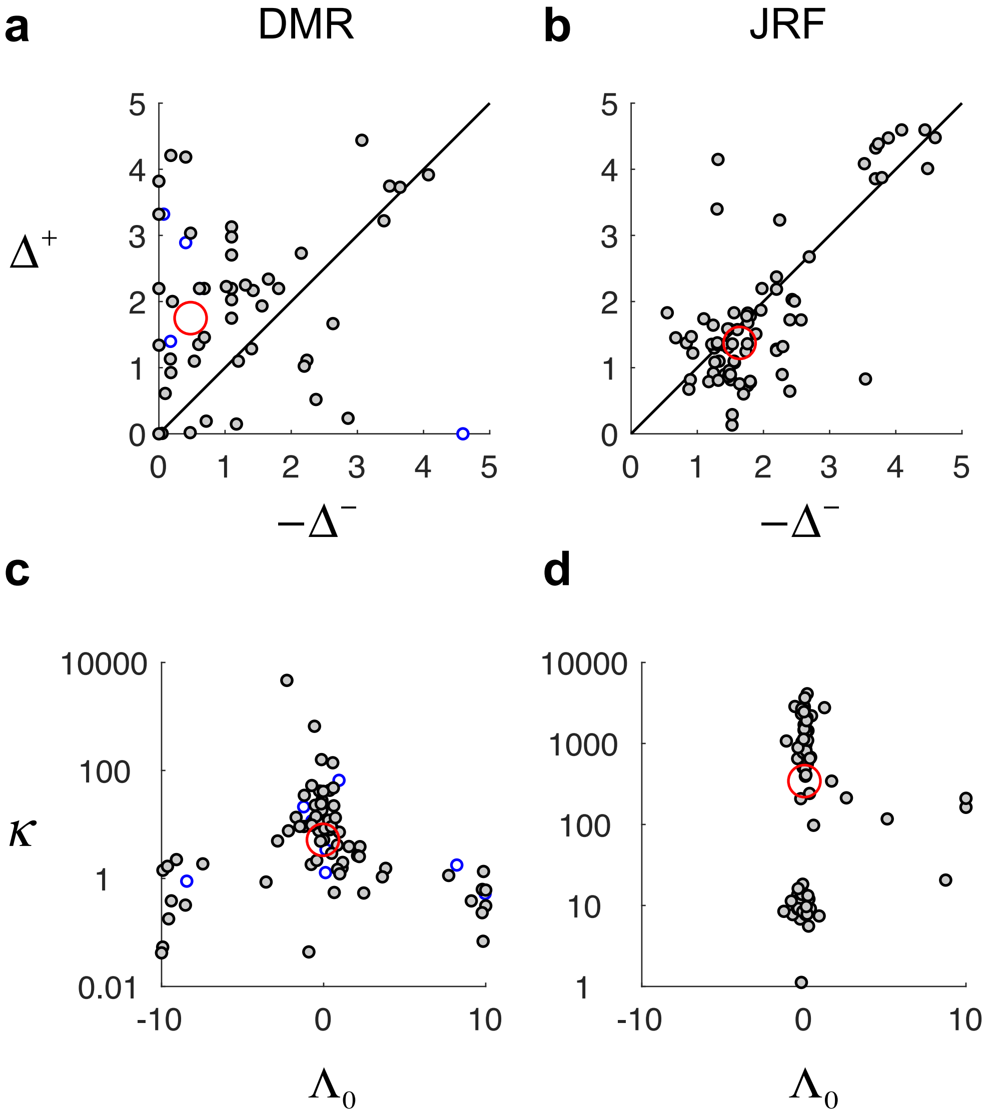


**Figure S7. The observed bounds and variance compensation parameters of BLO for individual participants.** Each small circle is for one participant, gray for Experiment JD, blue for Gonzalez and Wu (1999). The large red circle indicates the median across participants **a**. $\Delta^+$ versus $-\Delta^-$ for DMR. **b**. $\Delta^+$ versus $-\Delta^-$ for JRF. **c**. $\kappa$ versus $\Lambda_0$ for DMR. (2/85 data points are outside the plot range.) **d**. $\kappa$ versus $\Lambda_0$ for JRF. The value of $\kappa$ is plotted in log scale.

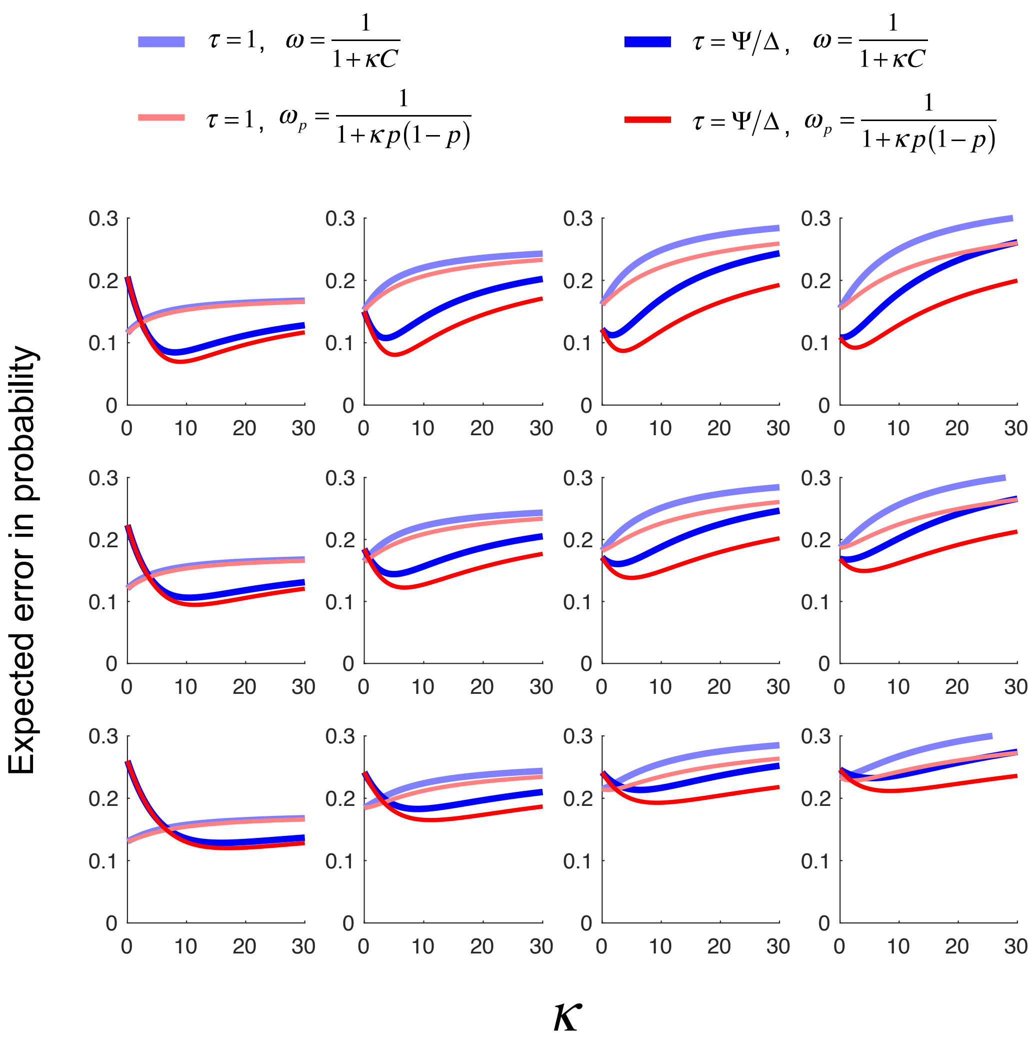


**Figure S8. Simulation results for variance compensation:** $\tau = 1$ **(lighter curves) versus** $\tau = \Psi/\Delta$ **(darker curves); constant** $\omega$ **(blue curves) versus** $\omega_p = \frac{1}{1+\kappa p(1-p)}$ **(red curves)**. To facilitate the comparison between $\omega_p$ and $\omega$, we write $\omega = \frac{1}{1+\kappa C}$, where $C$ denotes the median of *p(1–p)*. Expected error in probability is plotted as a function of $\kappa$. Each column is for one prior distribution of *p*. All distributions of objective log-odds $\lambda(p)$ are Gaussian distributions, whose standard deviations are respectively 0.8, 1.4, 2.0, 2.6, from left to right. Each row is for one noise level, from top to bottom corresponding to the 25%, 50%, and 75% percentile of Thurstone scale $\sigma_\Psi$ estimated from the 75 participants of our DMR experiment (see Supplement 11). In all the panels, the minimum expected error for $\tau = \Psi/\Delta$ is smaller than the minimum expected error for $\tau = 1$, and the minimum expected error for $\omega_p$ is smaller than the minimum expected error for constant $\omega$.

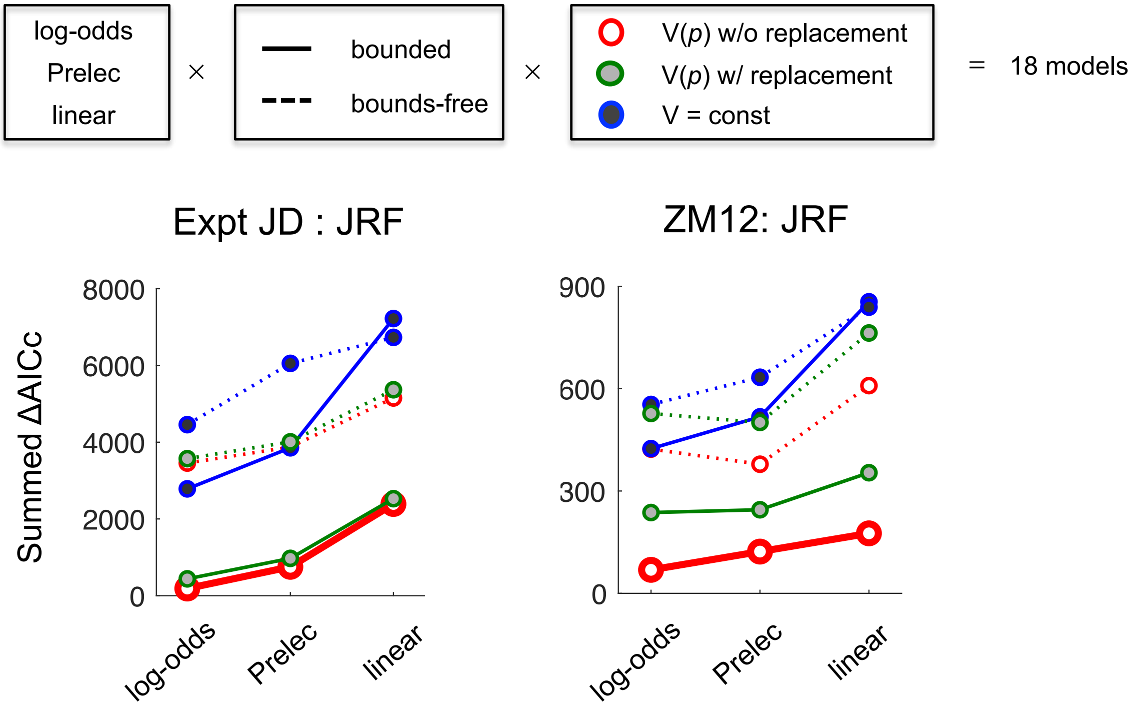


**Figure S9. Results of factorial model comparison for JRF including models assuming sampling with replacement.** Besides the 12 models presented in Figure 3, we considered 6 additional models whose assumption for variance compensation was based on sampling with replacement from the dot arrays (the BLO model presented in the main text assumes sampling without replacement). The summed ΔAICc across participants is plotted for each model, separately for the JRF tasks of Experiment JD (**a.** 75 participants) and Zhang and Maloney's (2012) Experiment 1 (**b.** 11 participants). Lower values of ΔAICc indicate better fits. Models with the assumption of sampling with replacement performed worse than those of sampling without replacement. Among the 18 models, the BLO model reported in the main text was still the best model.

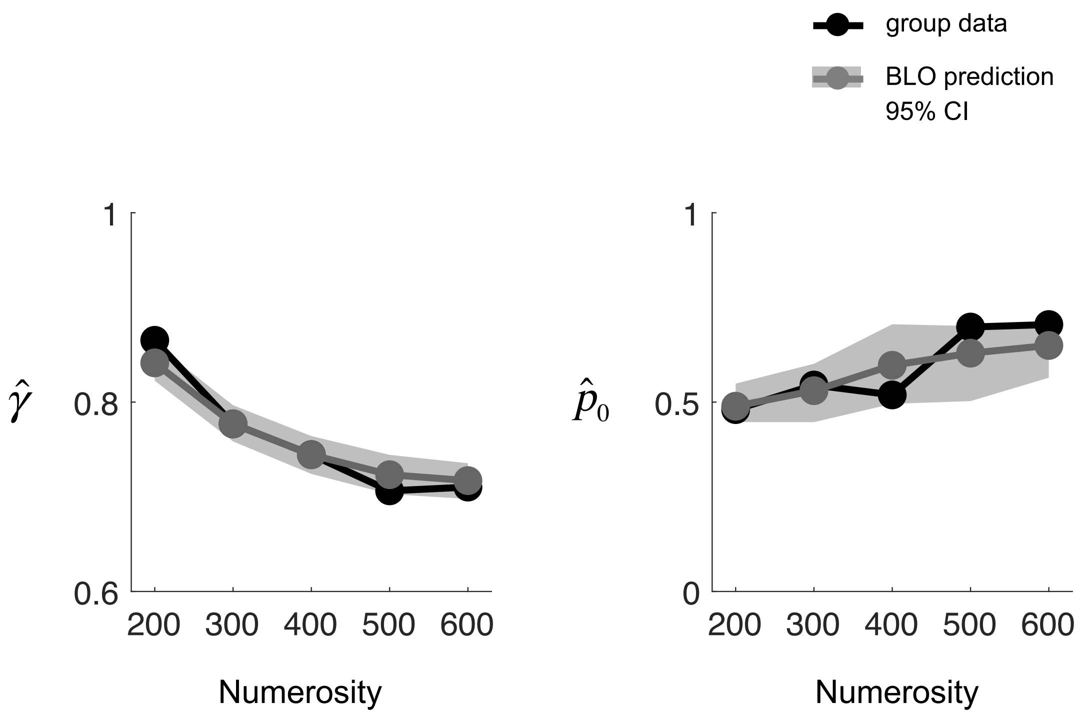


**Figure S10. Numerosity effect in Zhang and Maloney's (2012): data versus BLO prediction.** Left and right panels are respectively for $\hat{\gamma}$ and $\hat{p}_0$. The black line with dots denotes mean data across participants. The gray line with dots denotes the mean predicted by BLO and the gray shadow denotes its 95% confidence interval. BLO predictions agreed well with the data of Zhang and Maloney's (2012) Experiment 1, capturing the (significant) declining trend in the observed $\hat{\gamma}$ and the (marginally significant) rising trend in the observed $\hat{p}_0$.
